# diffcyt: Differential discovery in high-dimensional cytometry via high-resolution clustering

**DOI:** 10.1101/349738

**Authors:** Lukas M. Weber, Malgorzata Nowicka, Charlotte Soneson, Mark D. Robinson

## Abstract

High-dimensional flow and mass cytometry allow cell types and states to be characterized in great detail by measuring expression levels of more than 40 targeted protein markers per cell at the single-cell level. However, data analysis can be difficult, due to the large size and dimensionality of datasets as well as limitations of existing computational methods. Here, we present *diffcyt*, a new computational framework for differential discovery analyses in high-dimensional cytometry data, based on a combination of high-resolution clustering and empirical Bayes moderated tests adapted from transcriptomics. Our approach provides improved statistical performance, including for rare cell populations, along with flexible experimental designs and fast runtimes in an open-source framework.

## 2 Introduction

High-dimensional flow cytometry and mass cytometry (or CyTOF, for ‘cytometry by time-of-flight mass spectrometry’) characterize cell types and states by measuring expression levels of pre-defined sets of surface and intracellular proteins in individual cells, using antibodies tagged with either fluorochromes (flow cytometry) or heavy metal isotopes (mass cytometry). Modern flow cytometry systems allow simultaneous detection of more than 20 proteins per cell, in thousands of cells per second [1]. In mass cytometry, the use of metal tags significantly reduces signal interference due to spectral overlap and autofluorescence, enabling detection of more than 40 proteins per cell in hundreds of cells per second [1, 2]. Recently, further increases in the number of detected proteins have been demonstrated using oligonucleotide-tagged antibodies and single-cell sequencing [3]; this has also been combined with single-cell RNA sequencing on the same cells [4, 5].

The rapid increase in dimensionality has led to serious bottlenecks in data analysis. Traditional analysis by visual inspection of scatterplots (‘manual gating’) is unreliable and inefficient in high-dimensional data, does not scale readily, and cannot easily reveal unknown cell populations [1]. Significant efforts have been made to develop computationally guided or automated methods that do not suffer from these limitations. For example, unsupervised clustering algorithms are commonly used to define cell populations in one or more biological samples. Recent benchmarking studies have demonstrated that several clustering methods can accurately detect known cell populations in low-dimensional flow cytometry data [6], and both major and rare known cell populations in high-dimensional data [7]. A further benchmarking study comparing supervised methods for inferring cell populations associated with a censored continuous clinical variable demonstrated good performance for two methods using data of moderate dimensionality [8].

Several new methods have recently been developed for performing (partially) supervised analyses with the aim of inferring cell populations or states associated with an outcome variable in high-dimensional cytometry data, including *Citrus* [9], *CellCnn* [10], *cydar* [11], and a *classic* regression-based approach [12] (a similar regression-based approach was also recently described by [13]). However, these existing methods have a number of limitations. In particular: detected features from *Citrus* cannot be ranked by importance, and the ranking of detected cells from *CellCnn* cannot be interpreted in terms of statistical significance; rare cell populations are difficult to detect with *Citrus* and *cydar* (by contrast, *CellCnn* is optimized for analysis of rare populations); the response variable in the models for *Citrus* and *CellCnn* is the outcome variable, which makes it difficult to account for complex experimental designs; and *CellCnn* and *cydar* do not distinguish between ‘cell type’ and ‘cell state’ (e.g. functional) markers, which can make interpretation difficult.

Here, we present *diffcyt,* a new computational framework based on high-resolution unsupervised clustering together with supervised statistical analyses to detect cell populations or states associated with an outcome variable in high-dimensional cytometry data. The *diffcyt* methodology uses clustering to define cell populations, and empirical Bayes moderated tests adapted from transcriptomics for differential analysis. By default, our implementation uses the *FlowSOM* clustering algorithm [14], given its strong performance and fast runtimes [7]. For the differential analyses, we use methods from *edgeR* [15, 16], *limma* [17], and *voom* [18], which are widely used in the transcriptomics field; in addition, we include alternative methods adapted from the *classic* regression-based framework [12]. In principle, other high-resolution clustering algorithms or differential testing methods could also be substituted. Our methods consolidate several aspects of functionality from existing methods. Similar to *cydar* and the *classic* regression framework, our model specification uses the cytometry-measured features (cell population abundances or median expression of cell state markers within populations) as response variables, which enables analysis of complex experimental designs, including batch effects, paired designs, and continuous covariates. Linear contrasts enable testing of a wide range of hypotheses. Rare cell populations can easily be investigated, since the use of high-resolution clustering ensures that rare populations are unlikely to be merged into larger ones. In addition, as in *Citrus* and the *classic* regression framework, we optionally allow the user to split the set of protein markers into cell type and cell state markers. In this setup, cell type markers are used to define clusters representing cell populations, which are tested for differential abundance (DA); and median cell state marker signals per cluster are used to test for differential states (DS) within populations. We note that the underlying definitions of cell type and cell state can be challenging to apply to observed data, and may partially overlap. In general, cell type refers to relatively stable or permanent features of a cell’s identity, while cell state refers to transient features such as signaling or other functional states or the cell cycle [19–21]. In our view, providing the ability to maintain this distinction within the methodology greatly improves biological interpretability, since the results can be directly linked back to known cell types or populations of interest [12]. Finally, our methods have fast runtimes, enabling exploratory and interactive analyses.

## 3 Results

### 3.1 Overview and benchmarking strategy

Figure 1 provides a schematic overview of the *diffcyt* methodology (see Methods for further details), and Table 1 provides a summary of existing methods and their limitations. We demonstrate the performance of our methods using four benchmark datasets: two semi-simulated datasets *(AML-sim* and *BCR-XL-sim*) and two published experimental datasets *(Anti-PD-1* and *BCR-XL).* The semi-simulated datasets have been constructed by computationally introducing an artificial signal of interest (an *in silico* spike-in signal) into experimental data, thus reflecting the properties of real experimental data while also including a known ground truth that can be used to calculate statistical performance metrics. The experimental datasets, which do not contain a ground truth, are evaluated in qualitative terms. A complete description of all benchmark datasets is provided in Supplementary Note 1, and additional details on the comparisons with existing methods are included in Supplementary Note 2.

**Figure 1.**
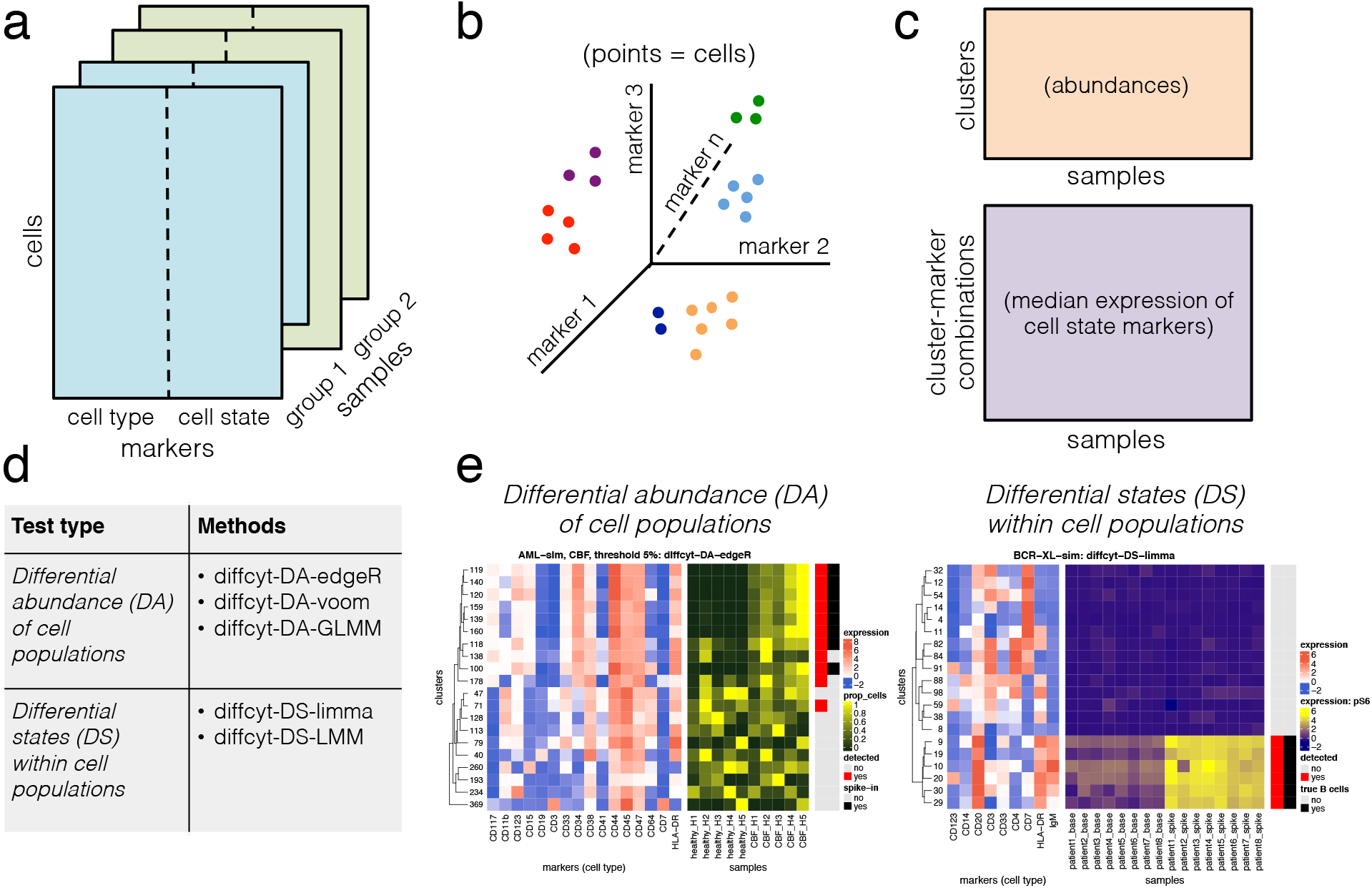
Schematic overview of diffcyt methodology. The *diffcyt* framework applies high-resolution clustering and empirical Bayes moderated tests for differential discovery analyses in high-dimensional cytometry data, (a) Input data are provided as tables of protein marker expression values per cell, one table per sample. Markers may be split into ‘cell type’ and ‘cell state’ categories; in the standard setup, cell type markers are used for clustering, (b) High-resolution clustering summarizes the data into a large number (e.g. 100-400) of clusters representing cell subsets, (c) Features are calculated at the cluster level, including cluster cell counts (abundances), and median expression of cell state markers within clusters, (d) Differential testing methods can be grouped into two types: differential abundance (DA) of cell populations, and differential states (DS) within cell populations. Results are returned in the form of adjusted p-values, allowing the identification of sets of significant detected clusters (DA tests) or cluster-marker combinations (DS tests), (e) Results are interpreted with the aid of visualizations, such as heatmaps. Example heatmaps show cluster phenotypes (expression profiles) and differential signal of interest (relative cluster abundances or expression of signaling marker pS6, by sample), with annotation for detected significant clusters or cluster-marker combinations (red) and true differential clusters or cluster-marker combinations (black). A detailed description of the *diffcyt* methodology is provided in Methods.

**Table 1.** Overview of existing methods and limitations. Overview of recently developed methods for performing differential analyses in high-dimensional cytometry data. For each method, a short description of the methodology and a summary of limitations are provided.

| Method | Short description | Limitations | Ref. |
| --- | --- | --- | --- |
| *Citrus* | Uses hierarchical clustering and regularized regression or classification models to select predictive features, such as cluster abundances or median expression of functional markers, that are associated with an outcome of interest. | • Detected features cannot be ranked by importance.<br>• Lasso-regularized models cannot easily detect multiple correlated features.<br>• Rare cell populations cannot easily be detected, due to minimum cluster size requirement and computational limitations.<br>• Response variable is the clinical outcome variable, which makes it difficult to account for complex experimental designs (including batch effects, paired designs, and continuous covariates). | [9] |
| *CellCnn* | Applies convolutional neural networks in a representation learning framework to detect rare cell populations associated with an outcome of interest. Designed specifically for detecting rare cell populations. | • Ranking of detected cells cannot be interpreted in terms of statistical significance.<br>• Interpretation of detected populations (referred to as filters) can be difficult, since they may be composed of multiple distinct cell populations.<br>• Response variable is the clinical outcome variable, which makes it difficult to account for complex experimental designs (including batch effects, paired designs, and continuous covariates).<br>• All protein markers are treated identically; there is no conceptual split between cell type and cell state (or functional) markers. | [10] |
| *cydar* | Assigns cells to overlapping hyperspheres in the high-dimensional space; tests for differential abundance between hyperspheres using moderated tests from *edgeR* [15, 16], while controlling the spatial false discovery rate among overlapping hyperspheres. | • Rare cell populations cannot easily be detected, due to their relatively small volume in the high-dimensional space.<br>• All protein markers are treated identically; there is no conceptual split between cell type and cell state (or functional) markers. | [11] |
| *classic regression-based approach* | Automated clustering using *FlowSOM* [14], followed by manual merging and annotation to define cell populations; differential testing of features such as population abundances or median expression of functional markers using generalized linear mixed models, linear mixed models, or linear models. | • Manual merging and annotation step requires expert biological knowledge, and can be time-consuming and subjective.<br>• When testing large numbers of clusters, e.g. to detect rare cell populations: loss of statistical power due to multiple testing penalty; no sharing of information across clusters. | [12] |

### 3.2 Improved performance for differential abundance tests

The *AML-sim* dataset evaluates performance for detecting differential abundance (DA) of rare cell populations (Figure 2). The dataset contains a spiked-in population of acute myeloid leukemia (AML) blast cells, in a comparison of 5 vs. 5 paired samples of otherwise healthy bone marrow mononuclear cells, which simulates the phenotype of minimal residual disease in AML patients (the data generation strategy is adapted from [10], and uses original data from [22]). The simulation was repeated for two subtypes of AML (cytogenetically normal, CN; and core binding factor translocation, CBF), and three thresholds of abundance for the spiked-in population (5%, 1%, and 0.1%). Figure 2(a) displays representative results for one subtype (CN) and one threshold (1%), for all *diffcyt* DA methods as well as *Citrus, CellCnn,* and *cydar* (complete results are included in Supplementary Figure 1). Methods *diffcyt-DA-edgeR, diffcyt-DA-voom,* and *CellCnn* give the best performance; the *diffcyt* results can also be interpreted as adjusted p-values, enabling a standard statistical framework where a list of significant detected clusters is determined by specifying a cutoff for the false discovery rate (FDR). *diffcyt-DA-GLMM* has inferior error control at the given FDR cutoffs, and reduced sensitivity at the highest spike-in threshold (5%). *Citrus* detects only a subset of the spiked-in cells, and *cydar* cannot reliably distinguish these rare populations. Figure 2(b) displays p-value distributions from an accompanying null simulation, where no true spike-in signal was included; the p-value distributions for the *diffcyt* methods are approximately uniform, indicating good error control and model fit (additional replicates are included in Supplementary Figure 2). Figure 2(c) illustrates the expression profiles (phenotypes) and relative abundances by sample for the detected and true differential clusters (additional heatmaps are included in Supplementary Figure 3). Figure 2(d) demonstrates the effect of varying the number of clusters across a broad range (between 9 and 1,600). Performance is reduced when there are too few clusters (due to merging of populations) or too many clusters (due to low power). The number of clusters is the main parameter choice in the *diffcyt* methods; an optimum is achieved around 400 clusters for this dataset (the remaining thresholds and condition are shown in Supplementary Figure 4).

**Figure 2.**
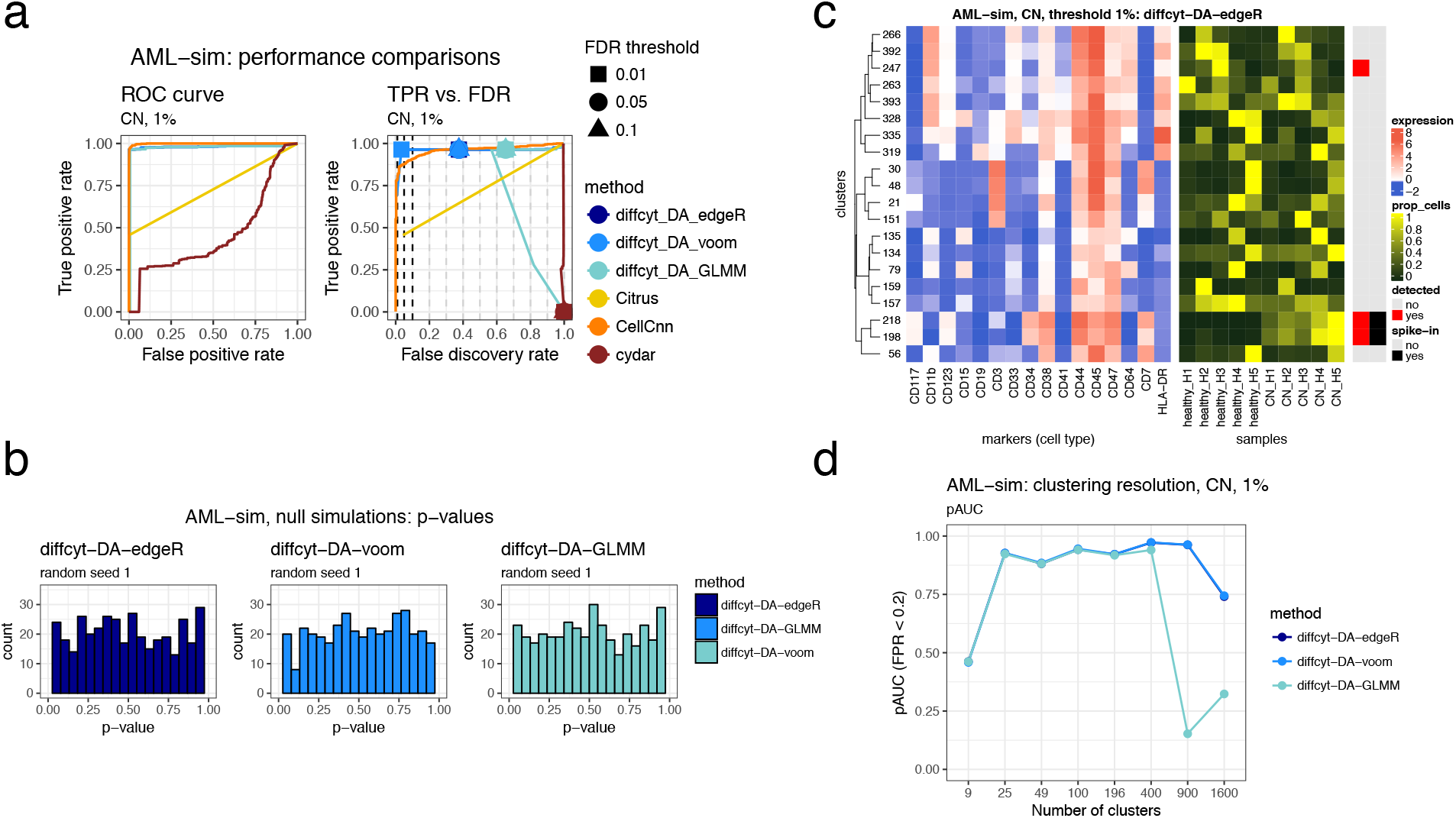
Benchmarking results for dataset AML-sim. (a) Performance metrics for dataset *AML-sim*, testing for differential abundance (DA) of cell populations. Panels show (i) receiver operating characteristic (ROC) curves, and (ii) true positive rate (TPR) vs. false discovery rate (FDR) (also indicating observed TPR and FDR at FDR cutoffs 1%, 5%, and 10%). Representative results for one condition (CN vs. healthy) and abundance threshold (1%) are shown (complete results for this dataset are included in Supplementary Figure 1). (b) Results for additional null simulations, where no true spike-in signal was included; p-value distributions are approximately uniform (additional replicates are included in Supplementary Figure 2). (c) Heatmap displaying phenotypes (expression profiles) of detected and true differential clusters, along with the signal of interest (relative cluster abundances, by sample), for method *diffcyt-DA-edgeR*. Expression values represent median arcsinh-transformed expression per cluster across all samples (left panel). Rows (clusters) are grouped by hierarchical clustering with Euclidean distance and average linkage; the heatmap shows the top 20 most highly significant clusters. Vertical annotation indicates detected significant clusters at 10% FDR (red) and clusters containing >50% true spiked-in cells (black). (Additional heatmaps are included in Supplementary Figure 3). (d) Results for varying clustering resolution (between 9 and 1,600 clusters); showing partial area under ROC curves (pAUC) for false positive rates (FPR) <0.2 (additional figures are included in Supplementary Figure 4). Performance metric plots generated using *iCOBRA* [23]; heatmaps generated using *ComplexHeatmap* [24].

Additional results provide further details on overall performance and robustness of the *diffcyt* DA methods. The top detected clusters represent high-precision subsets of the spiked-in population, confirming that the high-resolution clustering strategy has worked as intended (Supplementary Figure 5). Filtering clusters with low cell counts (using default parameters) did not remove any clusters from this dataset. An alternative implementation of the *diffcyt-DA-voom* method (using random effects for paired data) gives similar overall performance (Supplementary Figure 6). Using *FlowSOM* meta-clustering to generate 40 merged clusters instead of testing at high resolution worsens both error control and sensitivity (Supplementary Figure 7). The influence of random seeds used for the clustering and data generation procedures is greatest at the 0.1% threshold, as expected (Supplementary Figures 8-9). Similarly, additional simulations containing less distinct populations of interest (see Supplementary Note 1) show that reducing signal strength has a strong negative influence on performance at the 0.1% threshold (Supplementary Figure 10). Using smaller sample sizes (2 vs. 2) affects performance noticeably at the lower thresholds (Supplementary Figure 11). Finally, runtimes are fastest for methods *diffcyt-DA-edgeR* and *diffcyt-DA-voom* (Supplementary Figure 12).

### 3.3 Improved performance for differential state tests

The second dataset, *BCR-XL-sim,* evaluates performance for detecting differential states (DS) within cell populations (Figure 3). This dataset contains a spiked-in population of B cells stimulated with B cell receptor / Fc receptor cross-linker (BCR-XL), in a comparison of 8 vs. 8 paired samples of healthy peripheral blood mononuclear cells (original data sourced from [25]). The stimulated B cells have elevated expression of several signaling state markers, in particular phosphorylated ribosomal protein S6 (pS6); methods are evaluated by their ability to detect differential expression of pS6 within the population of B cells. Figure 3(a) summarizes performance for the *diffcyt* DS methods and the existing methods. The *diffcyt* methods give the best performance, with *diffcyt-DS-limma* having better error control. *Citrus* and *CellCnn* detect differential expression of pS6 for only a subset of the spiked-in cells, and *cydar* gives poor performance (likely due to ambiguity in assigning cells to overlapping hyperspheres in the high-dimensional space in order to calculate performance metrics). Figure 3(b) displays p-value distributions from a null simulation; p-values are approximately uniform across replicates, as previously (additional replicates are included in Supplementary Figure 13). Figure 3(c) displays expression profiles of detected and true differential clusters, along with expression by sample of the signaling marker pS6 (additional heatmaps are included in Supplementary Figure 14). Figure 3(d) demonstrates the effect of varying the number of clusters. Performance is reduced when there are too few or too many clusters; for this dataset, an optimum is observed across a broad range, including 100 clusters.

**Figure 3.**
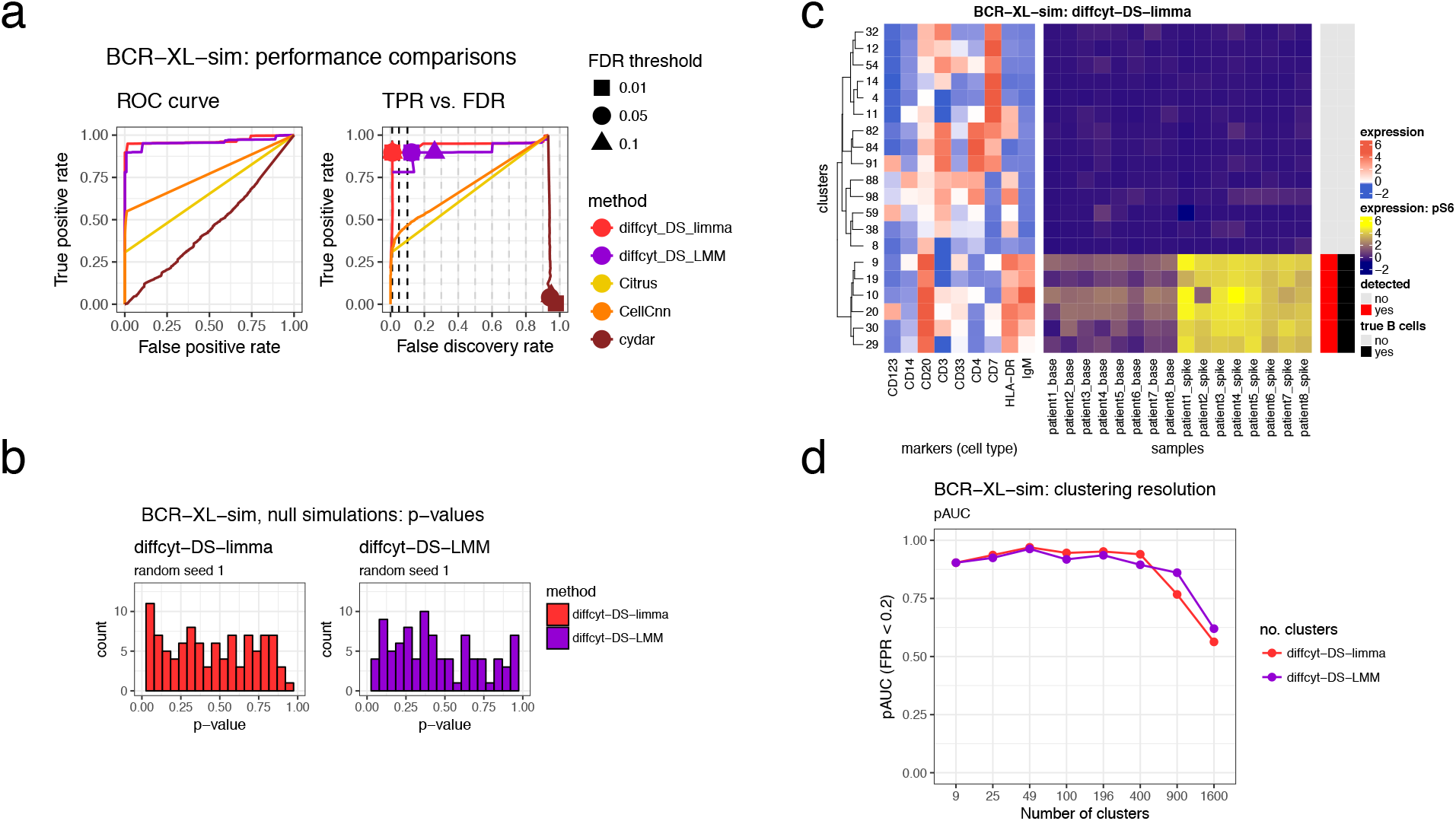
Benchmarking results for dataset BCR-XL-sim. (a) Performance metrics for dataset *BCR-XL-sim,* testing for differential states (DS) within cell populations. Panels show (i) receiver operating characteristic (ROC) curves, and (ii) true positive rate (TPR) vs. false discovery rate (FDR) (also indicating observed TPR and FDR at FDR cutoffs 1%, 5%, and 10%). (b) Results for additional null simulations, where no true spike-in signal was included; p-value distributions are approximately uniform (additional replicates are included in Supplementary Figure 13). (c) Heatmap displaying phenotypes (expression profiles) of detected and true differential clusters, along with the signal of interest (expression of signaling marker pS6, by sample), for method *diffcyt-DS-limma.* Expression values represent median arcsinh-transformed expression per cluster across all samples (left panel) or by individual samples (right panel). Rows (clusters) are grouped by hierarchical clustering with Euclidean distance and average linkage; the heatmap shows the top 20 most highly significant clusters. Vertical annotation indicates detected significant cluster-marker combinations at 10% FDR (red) and clusters containing >50% true spiked-in cells (black). (Additional heatmaps are included in Supplementary Figure 14). (d) Results for varying clustering resolution (between 9 and 1,600 clusters); showing partial area under ROC curves (pAUC) for false positive rates (FPR) <0.2. Performance metric plots generated using *iCOBRA* [23]; heatmaps generated using *ComplexHeatmap* [24].

As previously, the top detected clusters represent high-precision subsets of the population of interest (Supplementary Figure 15). Filtering with default parameters did not remove any clusters. To judge the benefit of splitting markers into cell type and cell state categories, we re-ran the analyses treating all markers as cell type (i.e. used for clustering), and using methods to test for DA instead of DS. This gave similar performance, but makes interpretation more difficult: since the methods test for DA of clusters defined using all markers in this case, the detected differential clusters may mix elements from canonical cell type and cell state phenotypes (Supplementary Figure 16). Alternative implementations of *diffcyt-DS-limma* (using random effects for paired data) and *diffcyt-DS-LMM* (using fixed effects for paired data) give similar performance overall (Supplementary Figure 17). For this dataset, using *FlowSOM* meta-clustering to merge clusters does not reduce performance (Supplementary Figure 18). Varying random seeds for the clustering and data generation procedures does not significantly affect performance (Supplementary Figures 19-20). Additional simulations containing less distinct populations of interest (see Supplementary Note 1) show deteriorating performance when the signal is reduced by 75% (Supplementary Figure 21). Using smaller sample sizes (4 vs. 4 and 2 vs. 2) worsens error control, especially for *diffcyt-DS-LMM* (Supplementary Figure 22). Runtimes are fastest for *diffcyt-DS-limma* (Supplementary Figure 23).

### 3.4 Successful recovery of known signals in experimental data

In order to demonstrate our methods on experimental data, we re-analyzed a dataset from a recent study using mass cytometry to characterize immune cell subsets in peripheral blood from melanoma patients treated with anti-PD-1 immunotherapy [26] *(Anti-PD-1* dataset; Figure 4). In this study, differential signals were detected for a number of cell populations, both in response to treatment and in baseline comparisons before treatment, between groups of patients classified as responders and non-responders to treatment. One key result was the identification of a small subpopulation of monocytes, with frequency in baseline samples (prior to treatment) strongly associated with responder status. The relatively rare frequency made this population difficult to detect; in addition, the dataset contained a strong batch effect due to sample acquisition on two different days [26]. Using method *diffcyt-DA-edgeR* to perform a differential comparison between baseline samples from the responder and non-responder patients (and taking into account the batch effect), we correctly identified three significant differentially abundant clusters (at an FDR cutoff of 10%) with phenotypes that closely matched the subpopulation of monocytes detected in the original study (CD14+ CD33+ HLA-DR^hi^ ICAM-1+ CD64+ CD141+ CD86+ CD11c+ CD38+ PD-L1+ CD11b+ monocytes) (clusters 317, 358, and 380; Figure 4(a)). One additional cluster with an unknown phenotype was also detected (cluster 308). The total abundance (combined cell counts) of the three matching clusters showed a clear differential signal between the two groups (Figure 4(b)). However, these results were sensitive to the choice of random seed for the clustering: in 5 additional runs using different random seeds, we detected between 0 and 4 significant differentially abundant clusters (at 10% FDR) per run; clusters matching the expected phenotype were detected in 4 out of the 5 runs (Supplementary Figure 24).

**Figure 4.**
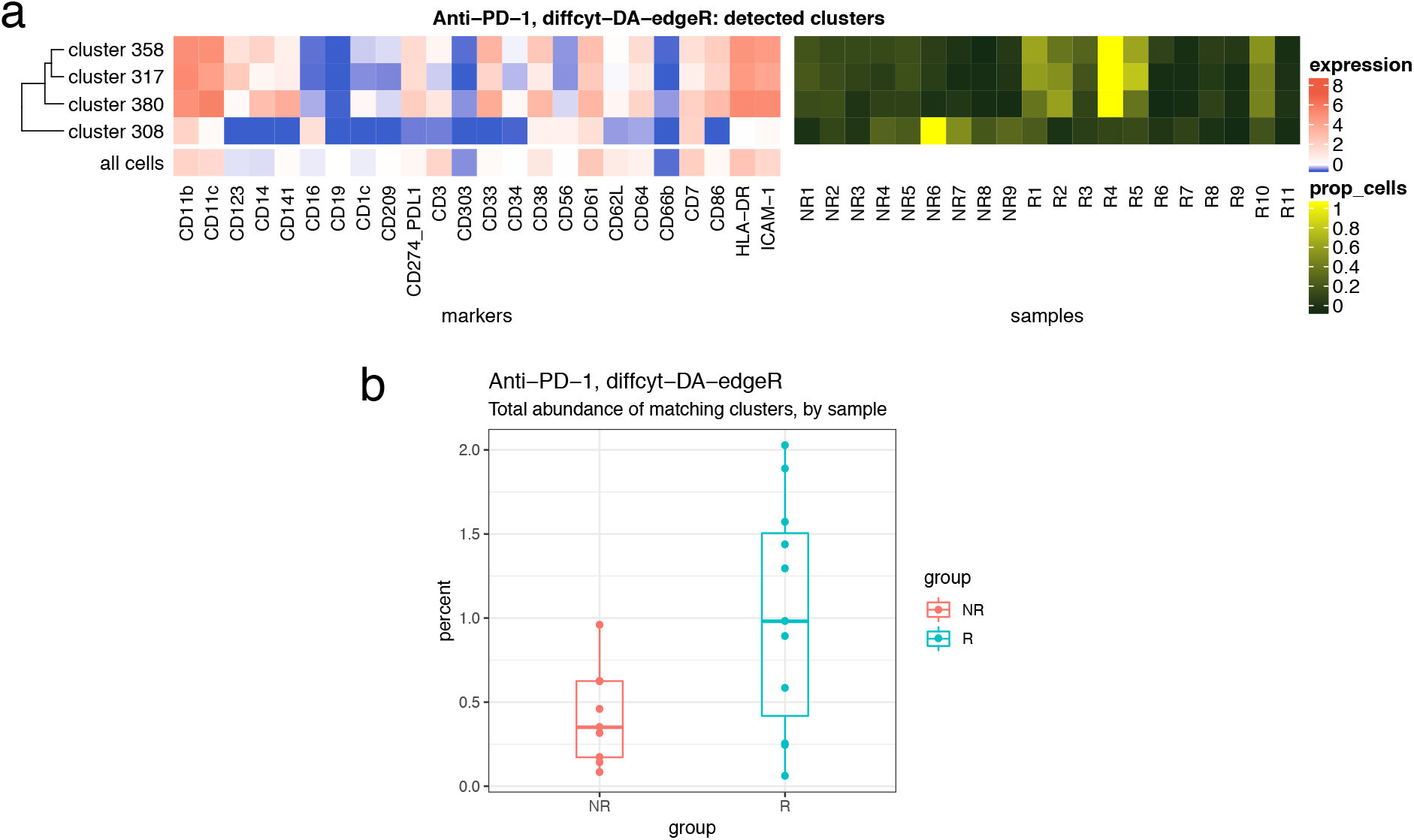
Results for experimental dataset Anti-PD-1. Results for re-analysis of experimental dataset *Anti-PD-1* using method *diffcyt-DA-edgeR*; testing for differential abundance (DA) of cell populations between baseline samples from responder and non-responder groups of patients. (a) Heatmap shows phenotype (median arcsinh-transformed marker expression profiles) of significant detected clusters at 10% false discovery rate (FDR), compared to all cells (left panel); and relative cluster abundances (proportion of cells per cluster, by sample) (right panel) for the detected clusters. Heatmap rows (clusters) are grouped by hierarchical clustering with Euclidean distance and average linkage. (b) Boxplot shows total abundance (combined number of cells) for the clusters matching the phenotype of interest (clusters 317, 358, and 380), by sample and group. Runtime was 32.0 seconds, on a 2014 MacBook Air laptop, 1.7 GHz processor, 8 GB memory, using a single processor core. NR = non-responders, R = responders.

For a second evaluation on experimental data, we re-analyzed the original (unmodified) data from the BCR-XL stimulation condition in [25] *(BCR-XL* dataset; Figure 5). This dataset contains strong differential signals for several signaling state markers in several cell populations, as previously described [12, 25]. Using method *diffcyt-DS-limma,* we reproduced several of the major known signals, including strong differential expression of: pS6, pPlcg2, pErk, and pAkt (elevated), and pNFkB (reduced, in BCR-XL stimulated condition) in B cells (identified by expression of CD20); pBtk and pNFkB in CD4+ T cells (identified by expression of CD3 and CD4); and pBtk, pNFkB, and pSlp76 in natural killer (NK) cells (identified by expression of CD7). Here, phenotypes can be identified either by marker expression profiles (Figure 5) or, alternatively, using reference population labels available for this dataset (Supplementary Figure 25).

**Figure 5.**
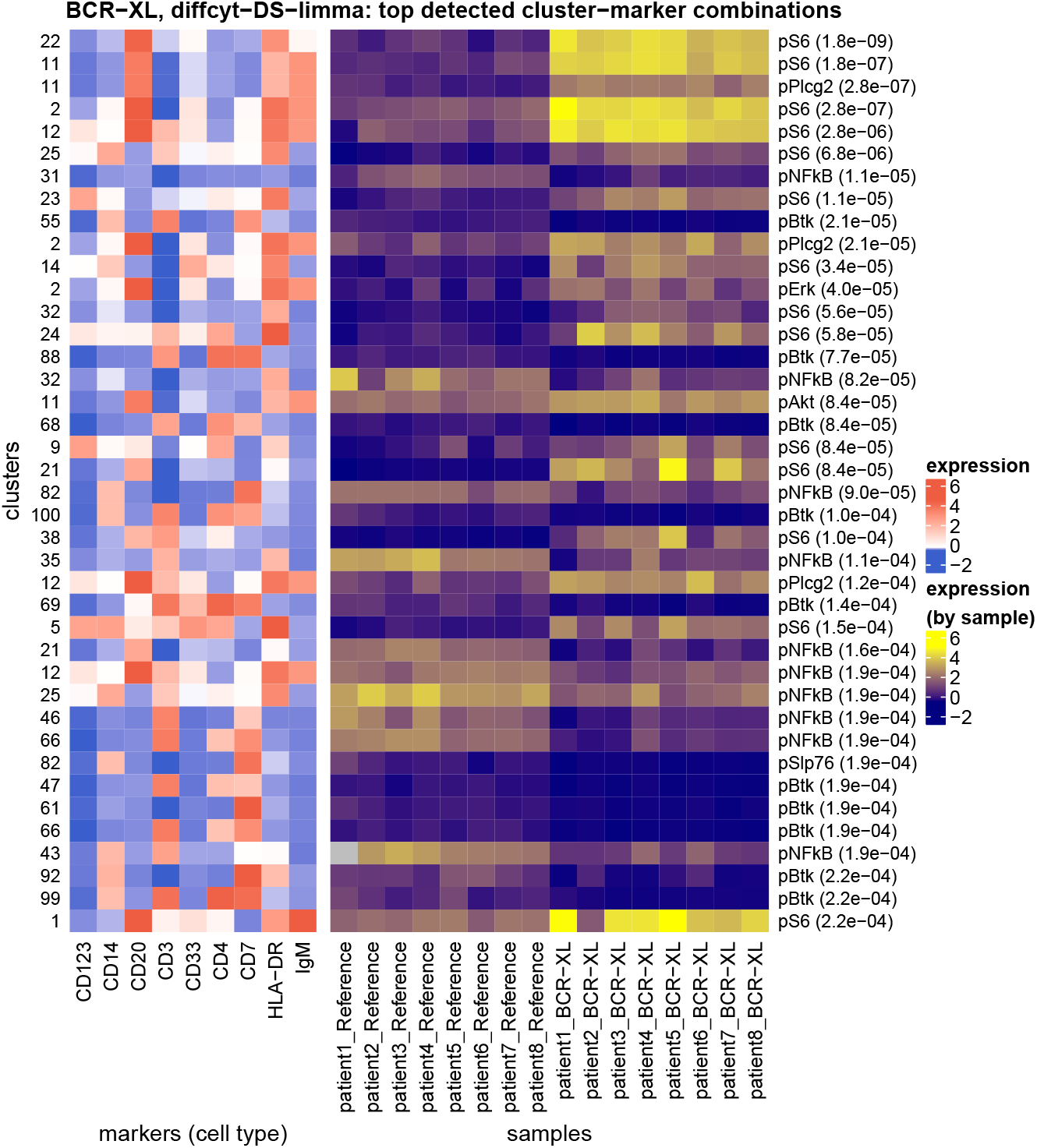
Results for experimental dataset BCR-XL. Results for re-analysis of experimental dataset *BCR-XL* using method *diffcyt-DS-limma*; testing for differential states (DS) within cell populations. Heatmap shows phenotypes (median arcsinh-transformed expression profiles for cell type markers) for the top 40 most highly significant detected cluster-marker combinations (left panel); and expression by sample for the cell state marker (signaling marker) in each detected cluster-marker combination (right panel). Rows (cluster-marker combinations) are ordered by decreasing adjusted p-values. Cell state marker names and adjusted p-values are displayed in right-hand-side row headings. Color scale for expression of cell type markers is normalized to 1st and 99th percentiles across all clusters and markers. Only the top 40 most highly significant detected cluster-marker combinations (out of 1,400 total) are shown, for easier visibility. Runtime was 15.0 seconds, on a 2014 MacBook Air laptop, 1.7 GHz processor, 8 GB memory, using a single processor core.

## 4 Discussion

We have presented a new computational framework for performing flexible differential discovery analyses in high-dimensional cytometry data. Our methods are designed for two related but distinct discovery tasks: detecting differentially abundant cell populations, including rare populations; and detecting differential expression of functional or other cell state markers within cell populations. Compared to existing approaches, our methods provide improved detection performance on semi-simulated benchmark datasets, along with fast runtimes. We have also successfully recovered known differential signals in re-analyses of two published experimental datasets, including differential abundance of a highly specific rare population. Our methods can account for complex experimental designs, including batch effects, paired designs, and continuous covariates. In addition, the set of protein markers may be split into cell type and cell state markers, facilitating biological interpretability. Visualizations such as heatmaps can be used to interpret the high-resolution clustering results (for example, to judge whether groups of clusters form larger populations, and to identify the phenotype of detected clusters). Methods *diffcyt-DA-edgeR* (for DA tests) and *diffcyt-DS-limma* (for DS tests) achieved the best performance and fastest runtimes overall (Figures 2–3); we recommend these as the default choices.

One limitation of our framework is that groups of similar clusters cannot be automatically merged into larger cell populations with a consistent phenotype. For example, the clear group of detected clusters in Figure 3(c) would ideally be merged into a single population representing B cells. However, this is a difficult computational problem, since the optimal resolution depends on the biological setting, and any automatic merging must avoid merging rare cell populations into larger ones. Our high-resolution clustering approach instead provides a tractable ‘middle ground’ between discrete clustering and a continuum of cell populations; we return results directly at the level of high-resolution clusters, and let the user interpret them via visualizations. A related issue concerns the identification of cell population phenotypes: our approach relies on visualizations and manual annotation of populations, which necessarily involves some subjectivity. Recently, several new methods have been published for automated labeling of cell populations [27], identification of simplified gating strategies to describe cell populations of interest [28, 29], or to compare cluster phenotypes [30]. These methods could be integrated within our framework to interpret detected differential clusters in a more automated manner. Similarly, clustering algorithms that generate biologically interpretable clusters could be used to improve interpretability [31].

A further limitation relates to batch effects: in datasets with strong batch effects, the high-resolution clustering may separate across batches, making it more difficult to distinguish the signal of interest. Aligning cell populations across batches is an active area of research in single-cell analysis (e.g. [32–35]); ideally, these methods will be integrated with frameworks for downstream differential analyses. Another issue concerns our strategy of summarizing cell state marker signals into median values. This strategy has advantages of simplicity, ease of interpretation, and fast runtimes. However, some information is necessarily lost, especially for markers with multi-modal distributions; good frameworks for flexible comparisons of full distributions are currently lacking. Additionally, splitting markers into groups representing cell type and cell state may be seen as a disadvantage in applications where this distinction is not clear. However, this step is optional: it is possible to run our methods using all markers for clustering (i.e. treating all markers as cell type) and testing for differential abundance (Supplementary Figure 16). For well-characterized immune populations, standard cell type markers may be found in the literature (e.g. [36]) or by consulting the websites of commercial antibody suppliers (e.g. BioLegend, Miltenyi Biotec, or Bio-Rad). Methods are also available to automatically group markers [12, 22, 37], although these should be used with care to ensure that cell population definitions are biologically plausible. For markers with subtle shifts (e.g. cytokines), assigning these as cell state markers and applying DS tests may fail to detect the differential signal; in this case, cluster labels may be exported to facilitate alternative analysis strategies (e.g. visualizations using *CytoRSuite* [38], *iSEE* [39], *OpenCyto* [40], or commercial software such as *FlowJo*).

The main user parameter in our methods is the number of clusters. The optimal value depends on several factors, including the size of the dataset (number of cells and samples), the expected relative abundances of cell populations of interest, and the number of markers used to define cell populations. The number of clusters determines the number of statistical tests, and affects power through the multiple testing penalty and the counts per cluster.

We recommend higher numbers of clusters when rare cell populations are of interest (for example, we used 400 clusters for the *AML-sim* dataset, and 100 clusters for the *BCR-XL-sim* dataset). Ultimately, this is a subjective choice for the user, which may also be explored interactively: e.g. by trying several different resolutions, and judging the interpretability of the results using visualizations or by calculating cluster separation metrics (e.g. average silhouette width). However, in our evaluations, good results were obtained over a range of resolutions (Figures 2(d) and 3(d)). Most computational methods include one or more parameters that can be adjusted by the user; in our view, one of the advantages of our approach is that the number of clusters is an intuitive parameter, with values that can be easily interpreted.

In general, we note that our methods are designed for ‘discovery’ analyses: all results should be explored and interpreted using visualizations, and any generated hypotheses must ultimately be validated with targeted confirmatory experiments. Our methods are implemented in the open-source R package *diffcyt,* available from Bioconductor (http://bioconductor.org/packages/diffcyt). The package includes comprehensive documentation and code examples, including an extended workflow vignette. Code to reproduce all analyses and figures from our benchmarking evaluations is available from GitHub (https://github.com/lmweber/diffcyt-evaluations), and data files from the benchmarking datasets are available from FlowRepository (FR-FCM-ZYL8) [41], allowing other researchers to extend and build on our analyses.

## 5 Methods

### 5.1 Description of *diffcyt* methodology

The following sections provide a detailed description of the *diffcyt* methodology (see Figure 1 for a schematic overview).

### 5.2 Preprocessing

#### Data preparation

Input data is formatted into a Bioconductor *SummarizedExperiment* object containing a single matrix of protein expression values, with one row per cell, and one column per protein marker. Row meta-data contains sample IDs and group IDs, and column meta-data contains protein marker information. The *SummarizedExperiment* format enables easy subsetting of data and meta-data, as well as simplified interaction with other Bioconductor packages.

#### Marker information: ‘cell type’ and ‘cell state’ markers

The set of protein markers may be split into sets of ‘cell type’ and ‘cell state’ markers. This split enables the methodology to take advantage of existing biological knowledge, and facilitates interpretability. By default, cell type markers are used to define clusters representing cell populations (which are tested for differential abundance), and median cell state marker signals are used to test for differential states (e.g. signaling or other functional states) within populations. This allows the user to interpret the results in terms of cell populations defined by known cell type markers.

The grouping into cell type and cell state markers must be specified by the user, and is stored in the column meta-data of the *SummarizedExperiment* object. This grouping is an important design choice, which may be made based on prior biological knowledge or using data-driven methods. For an example of a data-driven method of marker ranking and selection, see [22] and [12].

#### Subsampling

Optionally, random subsampling can be used to select an equal number of cells from each sample. This can be useful when there are large differences in total numbers of cells per sample, since it ensures that samples with relatively large numbers of cells do not dominate the clustering. However, some information will necessarily be lost. Subsampling should generally not be used when rare cell populations are of interest, due to the significant loss of information if cells from the rare population are discarded.

#### Transformation

Expression values are transformed using an inverse hyperbolic sine *(arcsinh*) transform with adjustable *cofactor* parameter. Raw expression values (fluorescence intensities for flow cytometry, or randomized ion counts for mass cytometry) follow an approximately log-normal distribution; the *arcsinh* transform brings this closer to a normal distribution (or mixture of normal distributions), which improves clustering performance and allows positive and negative populations to be distinguished more clearly. The *arcsinh* transform behaves similarly to a log transform at high values, but is approximately linear near zero; so unlike the log, it can handle zeros or small negative values. The cofactor controls the width of the linear region. (Zero values and small negatives occur in mass cytometry data when no ions are detected in a given channel: negatives are due to background subtraction and randomization of integer count values, which are performed by default by the instrument software). Standard values for the cofactor are 5 for mass cytometry, and 150 for flow cytometry (see [2], Supplementary Figure S2).

#### Integration with *CATALYST* package

Alternatively, a pre-prepared *daFrame* object from the *CATALYST* R/Bioconductor package [42] can be used as the input for the *diffcyt* methods. The *CATALYST* package contains extensive functions for preprocessing, exploratory analysis, and visualization of mass cytometry data. If this option is used, preprocessing (and clustering) are done using *CATALYST.* This is particularly useful when *CATALYST* has already been used for exploratory analyses and visualizations; the *diffcyt* package can then be used to calculate differential tests. For more details, see the *diffcyt* and *CATALYST* Bioconductor package vignettes.

### 5.3 Clustering

The clustering step is the core of the *diffcyt* methodology. We use high-resolution clustering to group cells into a large number of small clusters representing cell populations or subsets, which can then be further analyzed by differential testing. High-resolution clustering (or over-clustering) helps ensure that small or rare cell populations are adequately separated from larger populations.

By default, we use the *FlowSOM* clustering algorithm [14] (available from Bioconductor) to generate the clusters, since we previously showed that *FlowSOM* gives very good clustering performance for high-dimensional cytometry data, for both major and rare cell populations, and is extremely fast [7]. However, we run *FlowSOM* without the final meta-clustering step, to help ensure that small or rare populations are not merged into larger populations, which is crucial for detecting differential abundance of extremely rare populations.

If markers have been split into sets of cell type and cell state markers, then (by default) the clustering is performed using cell type markers only.

### 5.4 Data features

After clustering, we calculate features summarizing the data at the cluster level: cluster cell counts or abundances (number of cells per cluster-sample combination), and median transformed marker expression values (per cluster-sample combination). The feature values are formatted as new *SummarizedExperiment* objects, where rows represent clusters or cluster-marker combinations, and columns represent samples. These feature values are then used as inputs for the differential testing.

### 5.5 Design matrices and model formulas

The models to be fitted are specified with a design matrix or model formula, depending on the differential testing method used. Design matrices consist of one row per sample, and columns containing predictor variables, including the outcome of interest (e.g. columns of indicator variables for group IDs, such as diseased and healthy) and any other covariates. Flexible experimental designs are possible: block IDs (e.g. patient IDs in a paired design), batch effects, and continuous covariates can be included in the design matrix; each of these terms will be included as fixed effects in the models. Alternatively, model formulas also provide the option to include block IDs as random intercept terms (instead of fixed effects). When testing for differential abundance, model formulas can also be used to include random intercept terms for each sample (known as ‘observation-level random effects’ or OLREs; see [12]), to account for overdispersion typically seen in high-dimensional cytometry data.

### 5.6 Contrasts

The comparison of interest for the differential tests is specified with a contrast matrix. The contrast matrix consists of one row per model coefficient (corresponding to columns from the design matrix), and a column specifying the comparison of interest (i.e. the combination of model coefficients that is assumed to equal zero under the null hypothesis). This system of combining a design matrix (or model formula) with an appropriate contrast matrix provides users with powerful options to investigate a wide range of possible hypotheses within flexible experimental design settings.

### 5.7 Tests for differential abundance (DA) of cell populations

#### diffcyt-DA-edgeR

The *diffcyt-DA-edgeR* method calculates tests for differential abundance of clusters using methodology from the *edgeR* package [15, 16]. This method uses *edgeR* to fit models and calculate moderated tests at the cluster level. The moderated tests improve power by sharing information on variability (i.e. variance across samples for a single cluster) across clusters. Note that by default, we use the option *trend.method = “none”* to estimate common dispersions (see *edgeR* User’s Guide, available from Bioconductor).

The input to the tests is a table of cluster cell counts. The experimental design is specified using a design matrix, which enables flexible experimental designs. The comparison of interest is specified using a contrast matrix. A filtering step removes clusters with very low cell counts across samples to improve power. Normalization for the total number of cells per sample (library sizes) is automatically performed by the *edgeR* functions. Optionally, normalization factors for composition effects can be calculated using the ‘trimmed mean of M-values’ (TMM) method from the *edgeR* package [43].

Differential test results are returned in the form of raw p-values and adjusted p-values (FDR) from the moderated tests, which can be used to rank the clusters by their evidence for differential abundance. The results are stored in a new *SummarizedExperiment* object.

#### diffcyt-DA-voom

The *diffcyt-DA-voom* method calculates tests for differential abundance of clusters using methodology from the *limma* package [17] and *voom* method [18]. This method uses *limma* to fit models and calculate moderated tests at the cluster level. The moderated tests improve power by sharing information on variability across clusters. Since count data (such as cluster cell counts) are often heteroscedastic, we use *voom* to transform the raw cluster cell counts and estimate observation-level precision weights in order to stabilize the mean-variance relationship.

The input to the tests is a table of cluster cell counts. The experimental design is specified using a design matrix, which enables flexible experimental designs. For paired designs, either fixed effects or random effects can be used; fixed effects are simpler, but random effects may improve power in datasets with unbalanced designs or very large numbers of samples. Random effects make use of the *limma duplicateCorrelation* methodology (note that this methodology does not allow multiple measures per sample; in this case, fixed effects should be used instead). The comparison of interest is specified using a contrast matrix. A filtering step removes clusters with very low cell counts across samples to improve power. Normalization for the total number of cells per sample (library sizes) is automatically performed by the *limma* and *voom* functions. Optionally, normalization factors for composition effects can be calculated using the ‘trimmed mean of M-values’ (TMM) method from the *edgeR* package [43].

Differential test results are returned in the form of raw p-values and adjusted p-values (FDR) from the moderated tests, which can be used to rank the clusters by their evidence for differential abundance. The results are stored in a new *SummarizedExperiment* object.

#### diffcyt-DA-GLMM

The *diffcyt-DA-GLMM* method calculates tests for differential abundance of clusters using the generalized linear mixed models (GLMM) methodology originally implemented by [12]. This method fits GLMMs for each cluster, and calculates differential tests separately for each cluster (i.e. one model per cluster). The response variables in the models are the cluster cell counts, which are assumed to follow a binomial distribution. Note that the original methodology from [12] has been modified here to make use of high-resolution clustering to enable rare cell populations to be investigated more easily. In addition, we do not attempt to manually merge clusters into canonical cell populations; results are instead reported directly at the high-resolution cluster level.

The input to the tests is a table of cluster cell counts. The experimental design is specified using a model formula, which enables flexible experimental designs. Blocking variables (e.g. for paired designs) can be included as either random intercept terms or fixed effect terms. For paired designs, we recommend using random intercept terms to improve statistical power (see [12]). Batch effects and continuous covariates are included as fixed effects. In addition, we include random intercept terms for each sample to account for overdispersion typically seen in high-dimensional cytometry count data. The sample-level random intercept terms are known as ‘observation-level random effects’ (OLREs; see [12]). The comparison of interest is specified using a contrast matrix. A filtering step removes clusters with very low cell counts across samples to improve power. Optionally, normalization factors for composition effects can be calculated using the ‘trimmed mean of M-values’ (TMM) method from the *edgeR* package [43].

Differential test results are returned in the form of raw p-values and adjusted p-values (FDR), which can be used to rank the clusters by their evidence for differential abundance. The results are stored in a new *SummarizedExperiment* object.

### 5.8 Tests for differential states (DS) within cell populations

#### diffcyt-DS-limma

The *diffcyt-DS-limma* method calculates tests for differential states within clusters using methodology from the *limma* package [17]. Clusters are defined using cell type markers, and cell states are defined using median transformed expression of cell state markers within clusters. This method uses *limma* to fit models and calculate moderated tests at the cluster level. The moderated tests improve power by sharing information on variability across clusters. Note that by default, we use the option *trend = TRUE* in the *limma eBayes* fitting function in order to stabilize the mean-variance relationship.

The input to the tests is a set of tables of median expression of each marker for each cluster-sample combination. The experimental design is specified using a design matrix, which enables flexible experimental designs. For paired designs, either fixed effects or random effects can be used; fixed effects are simpler, but random effects may improve power in datasets with unbalanced designs or very large numbers of samples. Random effects make use of the *limma duplicateCorrelation* methodology (note that this methodology does not allow multiple measures per sample; in this case, fixed effects should be used instead). The comparison of interest is specified using a contrast matrix. A filtering step removes clusters with very low cell counts across samples to improve power. If cluster cell counts are provided, these can be used to calculate precision weights (across all samples and clusters), allowing the *limma* model fitting functions to account for uncertainty due to the total number of cells per sample (library size normalization) and total number of cells per cluster.

Differential test results are returned in the form of raw p-values and adjusted p-values (FDR) from the moderated tests for each cluster-marker combination (for cell state markers). These can be used to rank the cluster-marker combinations by their evidence for differential states. The results are stored in a new *SummarizedExperiment* object.

#### diffcyt-DS-LMM

The *diffcyt-DS-LMM* method calculates tests for differential states within clusters using the linear mixed models (LMM) and linear models (LM) methodology originally implemented by [12]. Clusters are defined using cell type markers, and cell states are defined using median transformed expression of cell state markers within clusters. This method fits LMMs for each cluster-marker combination (for cell state markers), and calculates differential tests separately for each cluster-marker combination (i.e. one model per cluster-marker combination). The response variable in each model is the median arcsinh-transformed marker expression of the cell state marker, which is assumed to follow a normal distribution. Note that the original methodology from [12] has been modified here to make use of high-resolution clustering to enable rare cell populations to be investigated more easily. In addition, we do not attempt to manually merge clusters into canonical cell populations; results are instead reported directly at the high-resolution cluster level.

The input is a set of tables of median expression of each marker for each cluster-sample combination. The experimental design is specified using a model formula, which enables flexible experimental designs. Blocking variables (e.g. for paired designs) can be included as either random intercept terms or fixed effect terms. For paired designs, we recommend using random intercept terms to improve statistical power (see [12]). Batch effects and continuous covariates are included as fixed effects. If no random intercept terms are included in the model formula, model fitting is performed using a linear model (LM) instead of a LMM. The comparison of interest is specified using a contrast matrix. A filtering step removes clusters with very low cell counts across samples to improve power. Within each model, sample-level weights can be included for the number of cells per sample; these weights represent the relative uncertainty in calculating each median value. (Additional uncertainty exists due to variation in the total number of cells per cluster; however, it is not possible to account for this, since separate models are used for each cluster-marker combination.)

Differential test results are returned in the form of raw p-values and adjusted p-values (FDR) for each cluster-marker combination (for cell state markers). These can be used to rank the cluster-marker combinations by their evidence for differential states. The results are stored in a new *SummarizedExperiment* object.

### 5.9 Interpretation and visualization

The *diffcyt* methods return results in the form of adjusted p-values (FDR) at the level of high-resolution clusters, either for a given cluster (for DA tests) or cluster-marker combination (for DS tests).

Due to the high-resolution clustering strategy, detected differential cell populations may be split into several sub-clusters with similar phenotypes. For biological interpretation, it is often useful to group the high-resolution clusters into larger populations with a consistent phenotype. However, automatically aggregating clusters is a difficult computational task, since the optimal resolution depends on the biological setting. In particular, there is a risk of merging rare cell populations into larger populations. Therefore, we have adopted the approach of returning results directly at the high-resolution cluster level. These results can then be explored and interpreted using visualizations.

Detailed visualizations can be generated using plotting functions from the *CATALYST* R/Bioconductor package [42], which accepts output objects from *diffcyt.* Key visualizations include heatmaps showing the phenotype (marker expression profiles) of detected clusters together with the sample-level signal of interest (cluster abundance or median expression of cell state markers). Examples are provided in the *diffcyt* and *CATALYST* Bioconductor package vignettes.

### 5.10 Number of clusters

The number of clusters is the main user parameter choice in the *diffcyt* methods. In the default implementation using the *FlowSOM* algorithm for clustering, this can be specified with the two arguments *xdim* and *ydim* in the function *generate Clusters*. The total number of clusters is then *xdim * ydim.* (This format is required since *FlowSOM* arranges clusters in a two-dimensional self-organizing map grid.)

The default is 100 clusters (*xdim = 10, ydim = 10*), which we expect is sufficient for many datasets. In general, we recommend higher numbers of clusters for datasets where rare cell populations are of interest. In our benchmarking evaluations, we used 400 clusters for the *AML-sim* dataset, and 100 clusters for the *BCR-XL-sim* dataset. Ultimately, this is a subjective choice for the user, which will depend on the biological setting and questions of interest in a given dataset; strategies to determine an appropriate number may include interactive exploration of visualizations, and (if available) making use of manually gated populations as a reference.

### 5.11 Benchmark datasets

A complete description of the benchmark datasets used to evaluate the methods is provided in Supplementary Note 1.

### 5.12 Comparisons with existing methods

Additional details on the comparisons with existing methods are provided in Supplementary Note 2.

## 6 Code and software availability

The methods described in this paper are implemented in the open-source R package d*iffcyt,* which is freely available from Bioconductor at http://bioconductor.org/packages/diffcyt. The *diffcyt* package includes comprehensive help files for each function, as well as a package vignette demonstrating a complete example workflow. Code scripts to reproduce all performance evaluations and comparisons with existing methods, reproduce all data preparation and simulation steps, and generate all figures, are available from GitHub at https://github.com/lmweber/diffcyt-evaluations. The results and figures in this paper were generated using *diffcyt* version 1.3.0 (available from GitHub at https://github.com/lmweber/diffcyt/releases) and R version 3.5.0.

## 7 Data availability

Data files for all benchmark datasets are available in FCS format from FlowRepository [41] (repository ID: FR-FCM-ZYL8) at http://flowrepository.org/id/FR-FCM-ZYL8. The benchmark datasets can also be accessed in *SummarizedExperiment* and *flowSet* Bioconductor object formats through the *HDCytoData* Bioconductor package, available at http://bioconductor.org/packages/HDCytoData.

## 8 Supplementary Material

Supplementary Material is available online as a single PDF document, containing Supplementary Figures 1-25 (supplementary results), Supplementary Note 1 (details on benchmark datasets), and Supplementary Note 2 (details on comparisons with existing methods). This is intended as a reference for consultation by readers interested in additional details, including details required for reproducing or extending results.

## 9 Author contributions

LMW and MDR developed methods, designed analyses, and wrote the manuscript. LMW implemented methods and performed analyses. MN developed methods and assisted with interpretation. CS assisted with designing analyses and interpretation. All authors read and approved the final manuscript.

## 10 Acknowledgments

The authors thank Helena L. Crowell (University of Zurich) for feedback on the implementation of the *diffcyt* R package, and all members of the Robinson Lab at the University of Zurich for feedback on the methodology and benchmarking. LMW was supported by a Forschungskredit (Candoc) grant from the University of Zurich (FK-17-100). MDR acknowledges support from the University Research Priority Program Evolution in Action at the University of Zurich.

## References

[1] Saeys, Y., Van Gassen, S., and Lambrecht, B. N. (2016). Computational flow cytometry: helping to make sense of high-dimensional immunology data. Nature Reviews Immunology, 16:449–462.

[2] Bendall, S. C., Simonds, E. F., Qiu, P., Amir, E.-a. D., Krutzik, P. O., Finck, R., Bruggner, R. V., Melamed, R., Trejo, A., Ornatsky, O. I., Balderas, R. S., Plevritis, S. K., Sachs, K., Pe’er, D., Tanner, S. D., and Nolan, G. P. (2011). Single-Cell Mass Cytometry of Differential Immune and Drug Responses Across a Human Hematopoietic Continuum. Science, 332:687–696.

[3] Shahi, P., Kim, S. C., Haliburton, J. R., Gartner, Z. J., and Abate, A. R. (2017). Abseq: Ultrahigh-throughput single cell protein profiling with droplet microfluidic barcoding. Scientific Reports, 7(44447).

[4] Stoeckius, M., Hafemeister, C., Stephenson, W., Houck-Loomis, B., Chattopadhyay, P. K., Swerdlow, H., Satija, R., and Smibert, P. (2017). Simultaneous epitope and transcriptome measurement in single cells. Nature Methods, 14(9):865–868.

[5] Peterson, V. M., Zhang, K. X., Kumar, N., Wong, J., Li, L., Wilson, D. C., Moore, R., McClanahan, T. K., Sadekova, S., and Klappenbach, J. A. (2017). Multiplexed quantification of proteins and transcripts in single cells. Nature Biotechnology, 35(10):936–939.

[6] Aghaeepour, N., Finak, G., The FlowCAP Consortium, The DREAM Consortium, Hoos, H., Mosmann, T. R., Brinkman, R., Gottardo, R., and Scheuermann, R. H. (2013). Critical assessment of automated flow cytometry data analysis techniques. Nature Methods, 10(3):228–238.

[7] Weber, L. M. and Robinson, M. D. (2016). Comparison of Clustering Methods for High-Dimensional Single-Cell Flow and Mass Cytometry Data. Cytometry Part A, 89A:1084–1096.

[8] Aghaeepour, N., Chattopadhyay, P., Chikina, M., Dhaene, T., Van Gassen, S., Kursa, M., Lambrecht, B. N., Malek, M., McLachlan, G. J., Qian, Y., Qiu, P., Saeys, Y., Stanton, R., Tong, D., Vens, C., Walkowiak, S., Wang, K., Finak, G., Gottardo, R., Mosmann, T., Nolan, G. P., Scheuermann, R. H., and Brinkman, R. R. (2016). A Benchmark for Evaluation of Algorithms for Identification of Cellular Correlates of Clinical Outcomes. Cytometry Part A, 89A:16–21.

[9] Bruggner, R. V., Bodenmiller, B., Dill, D. L., Tibshirani, R. J., and Nolan, G. P. (2014). Automated identification of stratifying signatures in cellular subpopulations. Proceedings of the National Academy of Sciences of the United States of America, pages E2770–E2777.

[10] Arvaniti, E. and Claassen, M. (2017). Sensitive detection of rare disease-associated cell subsets via representation learning. Nature Communications, 8(14825):1–10.

[11] Lun, A. T. L., Richard, A. C., and Marioni, J. C. (2017). Testing for differential abundance in mass cytometry data. Nature Methods, 14(7):707–709.

[12] Nowicka, M., Krieg, C., Weber, L. M., Hartmann, F. J., Guglietta, S., Becher, B., Levesque, M. P., and Robinson, M. D. (2017). CyTOF workflow: differential discovery in high-throughput high-dimensional cytometry datasets. F1000Research, version 2.

[13] Fonseka, C. Y., Rao, D. A., Teslovich, N. C., Korsunsky, I., Hannes, S. K., Slowikowski, K., Gurish, M. F., Donlin, L. T., Lederer, J. A., Weinblatt, M. E., Massarotti, E. M., Coblyn, J. S., Helfgott, S. M., Todd, D. J., Bykerk, V. P., Karlson, E. W., Ermann, J., Lee, Y. C., Brenner, M. B., and Raychaudhuri, S. (2018). Mixed-effects association of single cells identifies an expanded effector CD4+ T cell subset in rheumatoid arthritis. Science Translational Medicine, 10:eaaq0305.

[14] Van Gassen, S., Callebaut, B., Van Helden, M. J., Lambrecht, B. N., Demeester, P., Dhaene, T., and Saeys, Y. (2015). FlowSOM: Using Self-Organizing Maps for Visualization and Interpretation of Cytometry Data. Cytometry Part A, 87A:636–645.

[15] Robinson, M. D., McCarthy, D. J., and Smyth, G. K. (2010). edgeR: a Bioconductor package for differential expression analysis of digital gene expression data. Bioinformatics, 26(1):139–140.

[16] McCarthy, D. J., Chen, Y., and Smyth, G. K. (2012). Differential expression analysis of multifactor RNA-Seq experiments with respect to biological variation. Nucleic Acids Research, 40(10):4288–4297.

[17] Ritchie, M. E., Phipson, B., Wu, D., Hu, Y., Law, C. W., Shi, W., and Smyth, G. K. (2015). limma powers differential expression analyses for RNA-sequencing and microarray studies. Nucleic Acids Research, 43(7):e47.

[18] Law, C. W., Chen, Y., Shi, W., and Smyth, G. K. (2014). voom: precision weights unlock linear model analysis tools for RNA-seq read counts. Genome Biology, 15:R29.

[19] Wagner, A., Regev, A., and Yosef, N. (2016). Revealing the vectors of cellular identity with single-cell genomics. Nature Biotechnology, 34(11):1145–1160.

[20] Regev, A., Teichmann, S. A., Lander, E. S., Amit, I., Benoist, C., Birney, E., Bodenmiller, B., Campbell, P., Carninci, P., Clatworthy, M., Clevers, H., Deplancke, B., Dunham, I., Eberwine, J., Eils, R., Enard, W., Farmer, A., Fugger, L., Göttgens, B., Hacohen, N., Haniffa, M., Hemberg, M., Kim, S., Klenerman, P., Kriegstein, A., Lein, E., Linnarsson, S., Lundberg, E., Lundeberg, J., Majumder, P., Marioni, J. C., Merad, M., Mhlanga, M., Nawijn, M., Netea, M., Nolan, G., Pe’er, D., Phillipakis, A., Ponting, C. P., Quake, S., Reik, W., Rozenblatt-Rosen, O., Sanes, J., Satija, R., Schumacher, T. N., Shalek, A., Shapiro, E., Sharma, P., Shin, J. W., Stegle, O., Stratton, M., Stubbington, M. J. T., Theis, F. J., Uhlen, M., Oudenaarden, A. V., Wagner, A., Watt, F., Weissman, J., Wold, B., Xavier, R., Yosef, N., and Human Cell Atlas Meeting Participants (2017). The Human Cell Atlas. eLIFE, 6(e27041):1–30.

[21] Zeng, H. and Sanes, J. R. (2017). Neuronal cell-type classification: challenges, opportunities and the path forward. Nature Reviews Neuroscience, 18:530–546.

[22] Levine, J. H., Simonds, E. F., Bendall, S. C., Davis, K. L., Amir, E.-a. D., Tadmor, M. D., Litvin, O., Fienberg, H. G., Jager, A., Zunder, E. R., Finck, R., Gedman, A. L., Radtke, I., Downing, J. R., Pe’er, D., and Nolan, G. P. (2015). Data-Driven Phenotypic Dissection of AML Reveals Progenitor-like Cells that Correlate with Prognosis. Cell, 162:184–197.

[23] Soneson, C. and Robinson, M. D. (2016). iCOBRA: open, reproducible, standardized and live method benchmarking. Nature Methods, 13(4):283.

[24] Gu, Z., Eils, R., and Schlesner, M. (2016). Complex heatmaps reveal patterns and correlations in multidimensional genomic data. Bioinformatics, 32(18):2847–2849.

[25] Bodenmiller, B., Zunder, E. R., Finck, R., Chen, T. J., Savig, E. S., Bruggner, R. V., Simonds, E. F., Bendall, S. C., Sachs, K., Krutzik, P. O., and Nolan, G. P. (2012). Multiplexed mass cytometry profiling of cellular states perturbed by small-molecule regulators. Nature Biotechnology, 30(9):858–867.

[26] Krieg, C., Nowicka, M., Guglietta, S., Schindler, S., Hartmann, F. J., Weber, L. M., Dummer, R., Robinson, M. D., Levesque, M. P., and Becher, B. (2018). High-dimensional single-cell analysis predicts response to anti-PD-1 immunotherapy. Nature Medicine, 24(2):144–153.

[27] Abdelaal, T., van Unen, V., Höllt, T., Koning, F., Reinders, M. J., and Mahfouz, A. (2019). Predicting cell populations in single cell mass cytometry data. Cytometry Part A.

[28] Becht, E., Simoni, Y., Coustan-Smith, E., Evrard, M., Cheng, Y., Ng, L. G., Campana, D., and Newell, E. W. (2018). Reverse-engineering flow-cytometry gating strategies for phenotypic labelling and high-performance cell sorting. Bioinformatics, 35(2):301–308.

[29] Aghaeepour, N., Simonds, E. F., Knapp, D. J. H. F., Bruggner, R. V., Sachs, K., Culos, A., Gherardini, P. F., Samusik, N., Fragiadakis, G. K., Bendall, S. C., Gaudilliere, B., Angst, M. S., Eaves, C. J., Weiss, W. A., Fantl, W. J., and Nolan, G. P. (2018). GateFinder: projection-based gating strategy optimization for flow and mass cytometry. Bioinformatics, 34(23):4131–4133.

[30] Platon, L., Pejoski, D., Gautreau, G., Targat, B., Le Grand, R., Beignon, A.-S., and Tchitchek, N. (2018). A computational approach for phenotypic comparisons of cell populations in high-dimensional cytometry data. Methods, 132:66–75.

[31] Commenges, D., Alkhassim, C., Gottardo, R., Hejblum, B., and Thiébaut, R. (2018). cytometree: A Binary Tree Algorithm for Automatic Gating in Cytometry Analysis. Cytometry Part A, 93A:1132–1140.

[32] Butler, A., Hoffman, P., Smibert, P., Papalexi, E., and Satija, R. (2018). Integrating single-cell transcriptomic data across different conditions, technologies, and species. Nature Biotechnology, 36:411–420.

[33] Haghverdi, L., Lun, A. T. L., Morgan, M. D., and Marioni, J. C. (2018). Batch effects in singlecell RNA-sequencing data are corrected by matching mutual nearest neighbors. Nature Biotechnology, 36(5):421–427.

[34] Orlova, D. Y., Meehan, S., Parks, D., Moore, W. A., Meehan, C., Zhao, Q., Ghosn, E. E. B., Herzenberg, L. A., and Walther, G. (2018). QFMatch: multidimensional flow and mass cytometry samples alignment. Scientific Reports, 8(3291):1–14.

[35] Li, Y. H., Li, D., Samusik, N., Wang, X., Guan, L., Nolan, G. P., and Wong, W. H. (2017). Scalable multi-sample single-cell data analysis by Partition-Assisted Clustering and Multiple Alignments of Networks. PLoS Computational Biology, 13(12):1–37.

[36] Engel, P., Boumsell, L., Balderas, R., Bensussan, A., Gattei, V., Horejsi, V., Jin, B.-Q., Malavasi, F., Mortari, F., Schwartz-Albiez, R., Stockinger, H., van Zelm, M. C., Zola, H., and Clark, G. (2015). CD nomenclature 2015: Human leukocyte differentiation antigen workshops as a driving force in immunology. The Journal of Immunology, 195(10):4555–4563.

[37] Diggins, K. E., Greenplate, A. R., Leelatian, N., Wogsland, C. E., and Irish, J. M. (2017). Characterizing cell subsets using marker enrichment modeling. Nature Methods, 14(3):275–278.

[38] Hammill, D. (2019). CytoRSuite. R package, version 0.9.9.

[39] Rue-Albrecht, K., Marini, F., Soneson, C., Lun, A. T. L. (2018). iSEE: Interactive SummarizedExperiment Explorer. F1000Research, 7:741.

[40] Finak, G., Frelinger, J., Jiang, W., Newell, E. W., Ramey, J., Davis, M. M., Kalams, S. A., De Rosa, S. C., Gottardo, R. (2014). OpenCyto: An open source infrastructure for scalable, robust, reproducible, and automated, end-to-end flow cytometry data analysis. PLoS Computational Biology, 10(8):e1003806.

[41] Spidlen, J., Breuer, K., Rosenberg, C., Kotecha, N., and Brinkman, R. R. (2012). FlowRepository: A resource of annotated flow cytometry datasets associated with peer-reviewed publications. Cytometry Part A, 81A:727–731.

[42] Chevrier, S., Crowell, H. L., Zanotelli, V. R. T., Engler, S., Robinson, M. D., and Bodenmiller, B. (2018). Compensation of signal spillover in suspension and imaging mass cytometry. Cell Systems, 6:612–620.

[43] Robinson, M. D., Oshlack, A. (2010) A scaling normalization method for differential expression analysis of RNA-seq data. Genome Biology, 11:R25.

