## Supplementary Material for "diffcyt: Differential discovery in high-dimensional cytometry via high-resolution clustering"

Lukas M. Weber[1,2], Malgorzata Nowicka[1,2,†], Charlotte Soneson[1,2,‡], Mark D. Robinson[1,2,*]

[1]Institute of Molecular Life Sciences, University of Zurich, Zurich, Switzerland
[2]SIB Swiss Institute of Bioinformatics, University of Zurich, Zurich, Switzerland

†Current address: F. Hoffmann-La Roche AG, Basel, Switzerland

‡Current address: Friedrich Miescher Institute for Biomedical Research and SIB Swiss Institute of Bioinformatics, Basel, Switzerland

*Corresponding author

March 28, 2019

### Supplementary Figures

#### 1.1 AML-sim: Differential abundance of rare cell populations

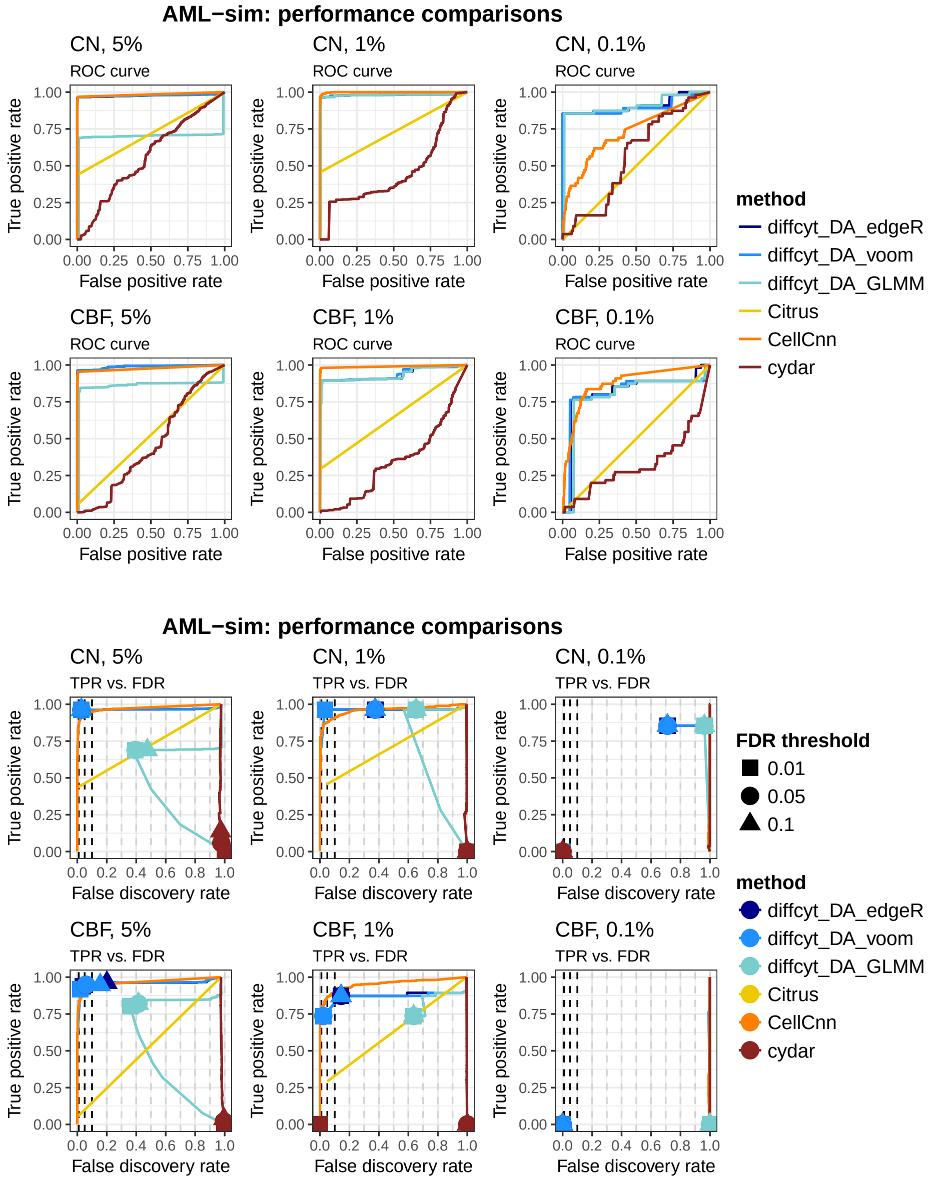

**Supplementary Figure 1. Performance metrics: AML-sim, all methods.** Results of main performance evaluations for `diffcyt` methods and comparisons with `Citrus`, `CellCnn`, and `cydar`; testing for differential abundance of rare cell populations. Panels display results for condition CN vs. healthy (rows 1 and 3) and condition CBF vs. healthy (rows 2 and 4), at three different thresholds of abundance for the rare cell population (5%, 1%, and 0.1%; by column). Panels show (i) receiver operating characteristic (ROC) curves, and (ii) true positive rate (TPR) vs. false discovery rate (FDR) curves (also showing observed TPR and FDR at FDR cutoffs 1%, 5%, and 10%). See section 3.4 for additional notes on the evaluation metrics for each method.

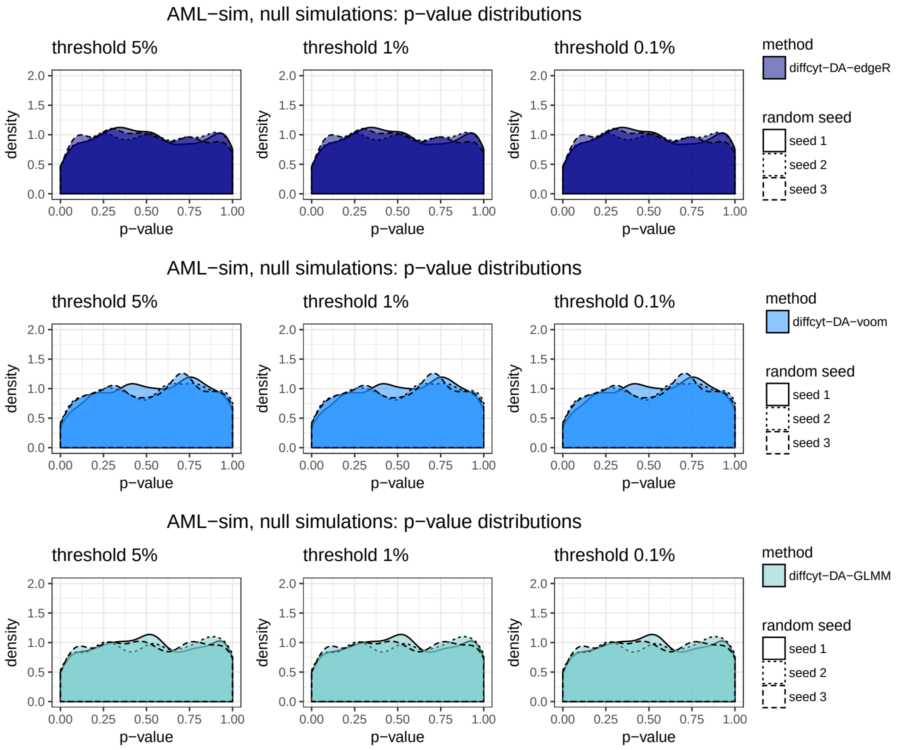

**Supplementary Figure 2. Null simulation p-values: AML-sim, diffcyt methods.** P-value distributions (densities) for `diffcyt` methods, 3 replicated null simulations; testing for differential abundance of rare cell populations. Panels display results for three different thresholds of abundance for the rare cell population (5%, 1%, and 0.1%; by column), for each method (by row and color). P-value distributions that are approximately uniform across replicates are consistent with data containing no true differential signal.

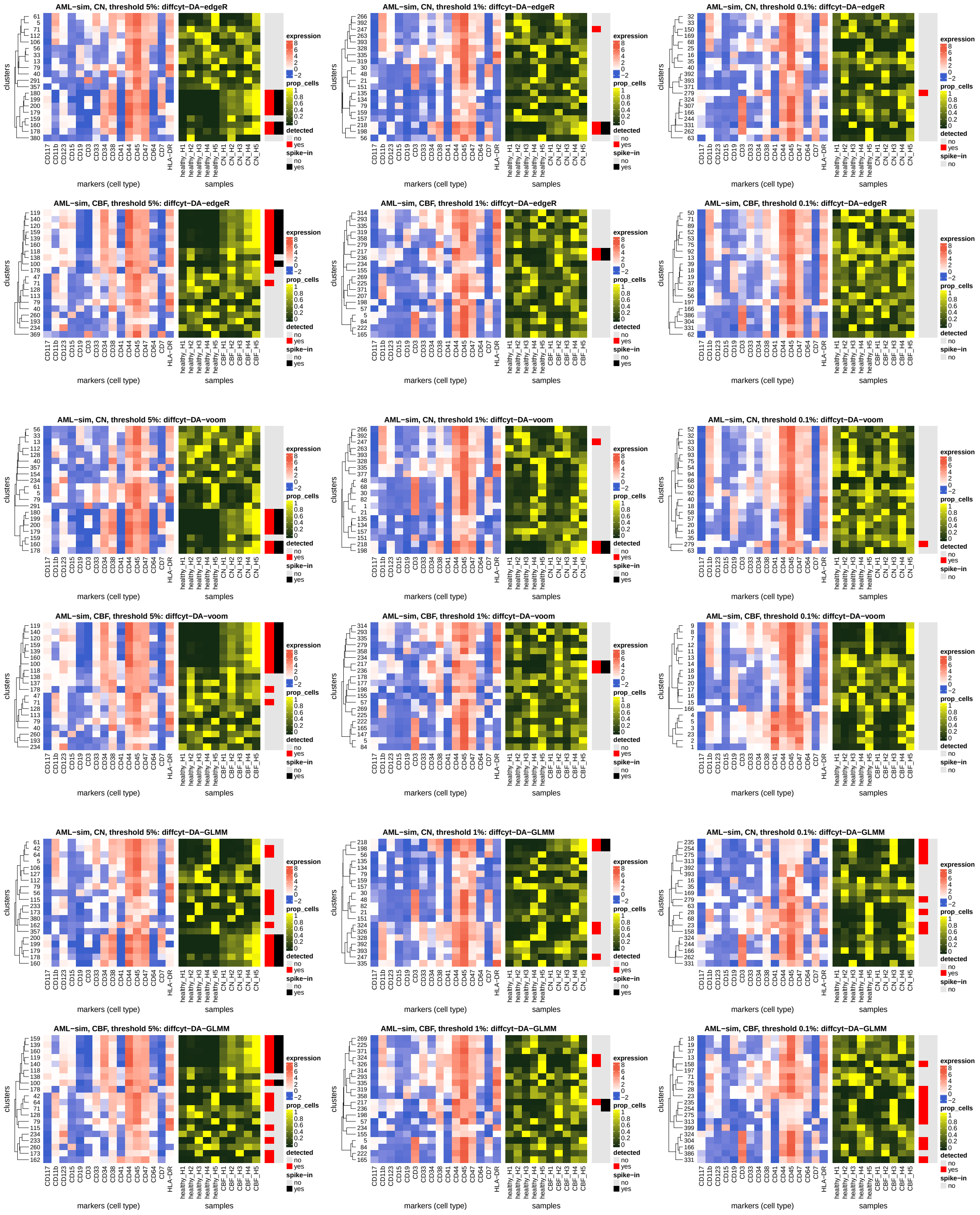

**Supplementary Figure 3. Heatmaps: AML-sim, diffcyt methods.** Heatmaps showing (i) phenotypes (median arcsinh-transformed expression profiles for cell type markers) (left panel) and (ii) relative cluster abundances (proportion of cells per cluster, by sample); testing for differential abundance of rare cell populations. Vertical annotation highlights detected significant clusters at 10% FDR (red) and clusters containing >50% true spiked-in cells (black). Color scale for expression is normalized to 1st and 99th percentiles across all clusters and cell type markers. Clusters (rows) are grouped using hierarchical clustering with Euclidean distance and average linkage. Each heatmap shows only the top 20 clusters as ranked by significance levels (out of 400 clusters total), for easier visibility of the top detected clusters. Panels show results for condition CN vs. healthy and condition CBF vs. healthy (arranged in rows), at three different thresholds of abundance for the rare cell population (5%, 1%, and 0.1%; arranged in columns), for each method (arranged by group).

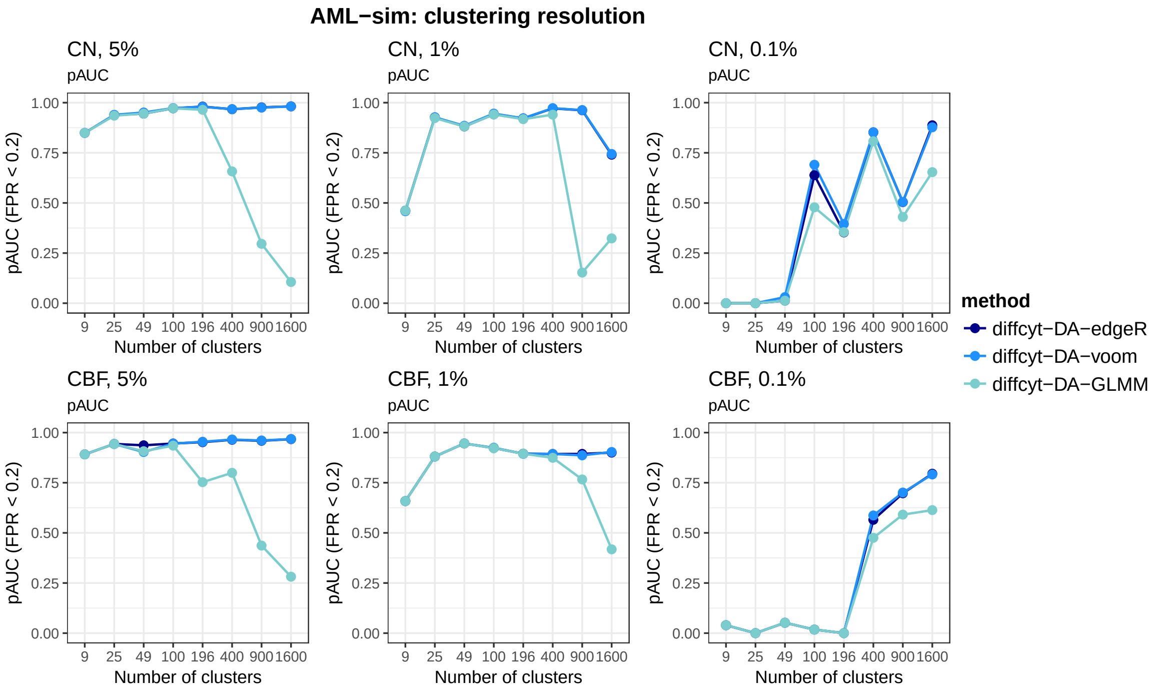

**Supplementary Figure 4. Performance metrics: AML-sim, diffcyt methods, varying clustering resolution.** Results of performance evaluations for `diffcyt` methods, varying clustering resolution; testing for differential abundance of rare cell populations. Plots show partial area under receiver operating characteristic (ROC) curves (pAUC) for false positive rates (FPR) $< 0.2$, for selected numbers of clusters. Panels display pAUC plots for condition CN vs. healthy (row 1) and condition CBF vs. healthy (row 2), at three different thresholds of abundance for the rare cell population (5%, 1%, and 0.1%; by column).

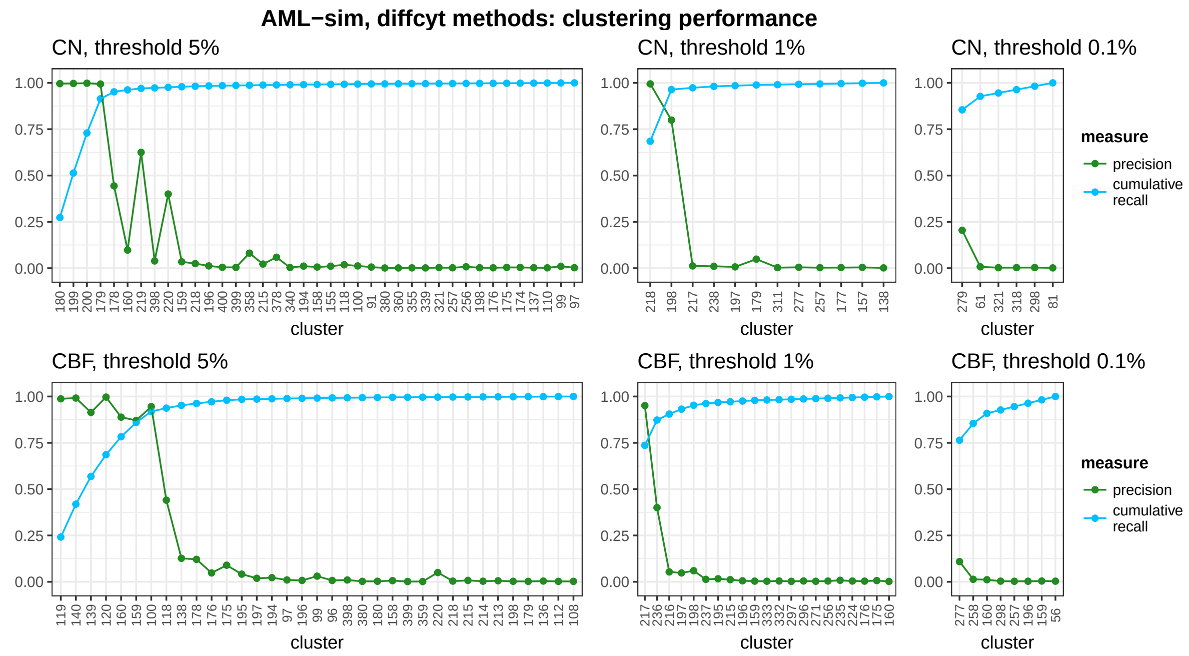

**Supplementary Figure 5. Clustering performance: AML-sim, diffcyt methods.** Clustering performance measures (precision, cumulative recall) for all clusters containing any true spiked-in cells (recall $> 0$); testing for differential abundance of rare cell populations. Clusters (horizontal axis) are ordered by recall. Note that the clustering step is the same for all `diffcyt` methods (`diffcyt-DA-edgeR`, `diffcyt-DA-voom`, and `diffcyt-DA-GLMM`). Panels show results for condition CN vs. healthy (row 1) and condition CBF vs. healthy (row 2), at three different thresholds of abundance for the rare cell population (5%, 1%, and 0.1%; by column).

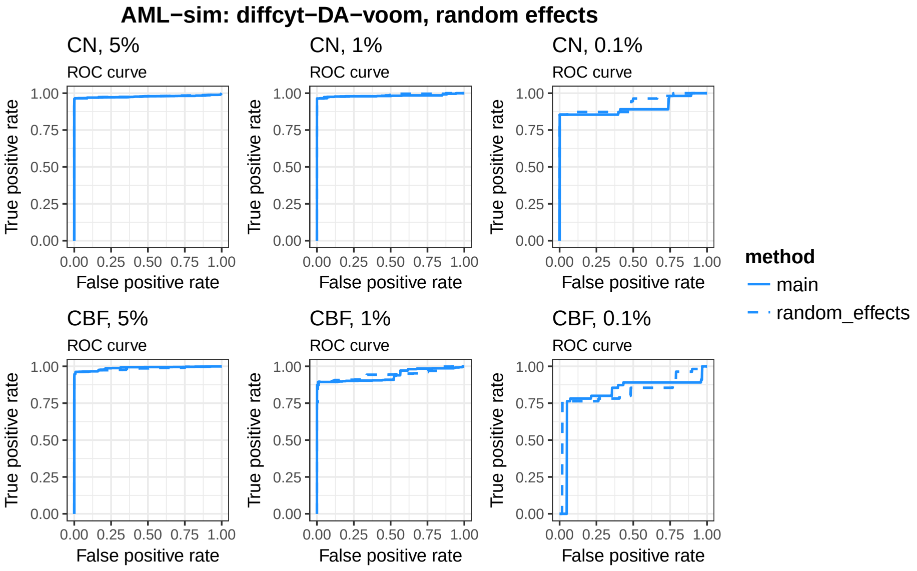

**Supplementary Figure 6. Performance metrics: AML-sim, diffcyt-DA-voom, random effects for patient IDs.** Results of performance evaluations for `diffcyt-DA-voom`, main results vs. using random effects instead of fixed effects for patient IDs (using `limma duplicateCorrelation` methodology); testing for differential abundance of rare cell populations. Panels display receiver operating characteristic (ROC) curves for condition CN vs. healthy (row 1) and condition CBF vs. healthy (row 2), at three different thresholds of abundance for the rare cell population (5%, 1%, and 0.1%; by column).

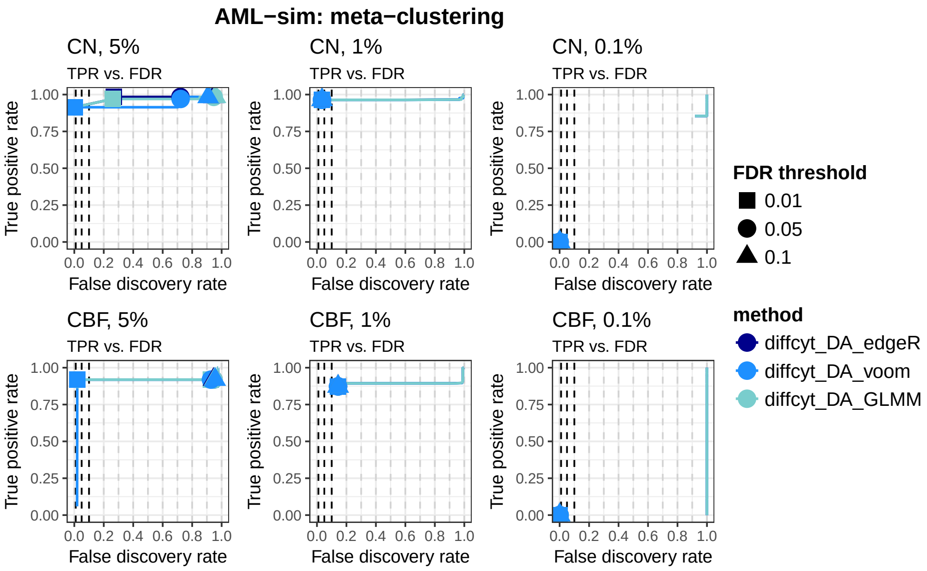

**Supplementary Figure 7. Performance metrics: AML-sim, diffcyt methods, using FlowSOM meta-clustering.** Results of performance evaluations for `diffcyt` methods, using 40 meta-clusters in FlowSOM clustering algorithm; testing for differential abundance of rare cell populations. Panels display true positive rate (TPR) vs. false discovery rate (FDR) curves (also showing observed TPR and FDR at FDR cutoffs 1%, 5%, and 10%) for condition CN vs. healthy (row 1) and condition CBF vs. healthy (row 2), at three different thresholds of abundance for the rare cell population (5%, 1%, and 0.1%; by column).

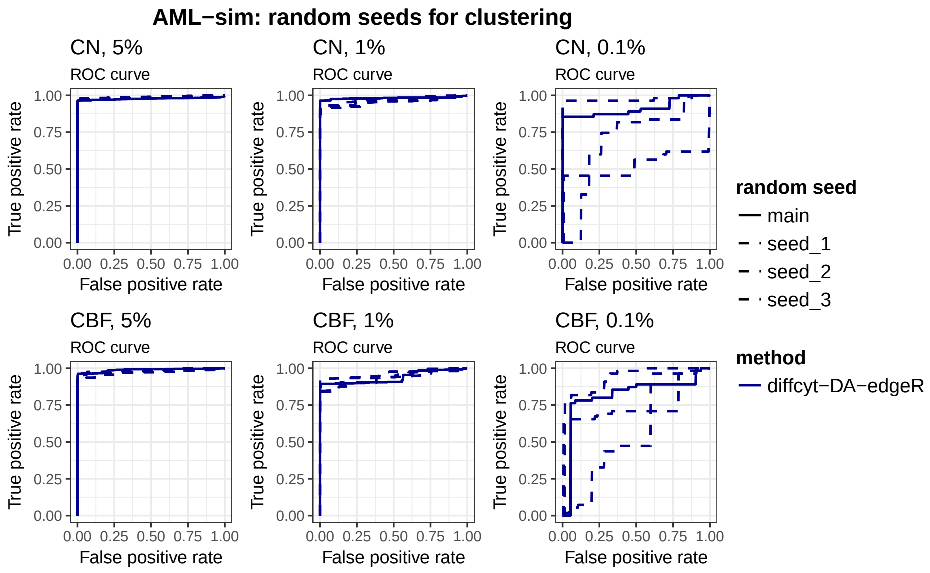

**Supplementary Figure 8. Performance metrics: AML-sim, diffcyt-DA-edgeR, varying random seeds for clustering.** Results of performance evaluations for `diffcyt-DA-edgeR`, main results and 3 additional replicates using varying random seeds for clustering step; testing for differential abundance of rare cell populations. Panels display receiver operating characteristic (ROC) curves for condition CN vs. healthy (row 1) and condition CBF vs. healthy (row 2), at three different thresholds of abundance for the rare cell population (5%, 1%, and 0.1%; by column). Results for methods `diffcyt-DA-voom` and `diffcyt-DA-GLMM` are approximately similar.

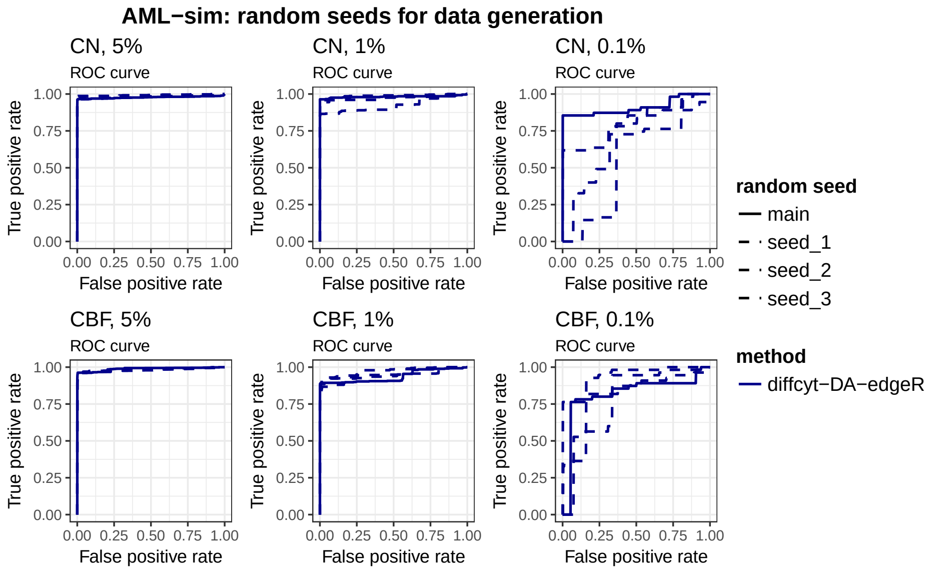

**Supplementary Figure 9. Performance metrics: AML-sim, diffcyt-DA-edgeR, varying random seeds for data generation.** Results of performance evaluations for `diffcyt-DA-edgeR`, main results and 3 additional replicates using varying random seeds for data generation; testing for differential abundance of rare cell populations. Panels display receiver operating characteristic (ROC) curves for condition CN vs. healthy (row 1) and condition CBF vs. healthy (row 2), at three different thresholds of abundance for the rare cell population (5%, 1%, and 0.1%; by column). Results for methods `diffcyt-DA-voom` and `diffcyt-DA-GLMM` are approximately similar.

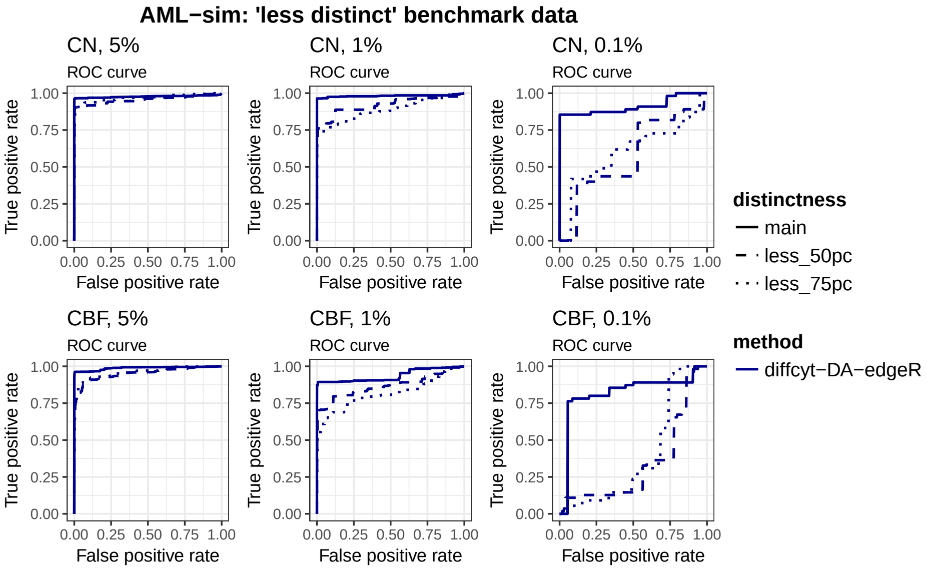

**Supplementary Figure 10. Performance metrics: AML-sim, diffcyt-DA-edgeR, 'less distinct' benchmark data.** Results of performance evaluations for `diffcyt-DA-edgeR`, main results and 50% and 75% 'less distinct' benchmark datasets; testing for differential abundance of rare cell populations. Panels display receiver operating characteristic (ROC) curves for condition CN vs. healthy (row 1) and condition CBF vs. healthy (row 2), at three different thresholds of abundance for the rare cell population (5%, 1%, and 0.1%; by column). Results for methods `diffcyt-DA-voom` and `diffcyt-DA-GLMM` are approximately similar.

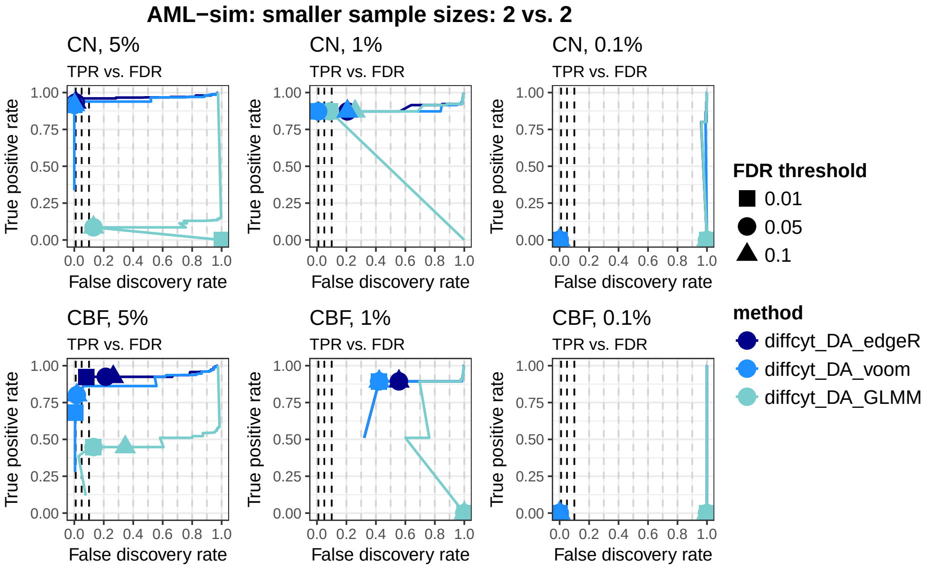

**Supplementary Figure 11. Performance metrics: AML-sim, diffcyt methods, smaller sample sizes.** Results of performance evaluations for `diffcyt` methods, using a subset of the full number of samples (2 vs. 2 samples); testing for differential abundance of rare cell populations. The full dataset contains 5 vs. 5 samples. Panels display true positive rate (TPR) vs. false discovery rate (FDR) curves (also showing observed TPR and FDR at FDR cutoffs 1%, 5%, and 10%) for condition CN vs. healthy (row 1) and condition CBF vs. healthy (row 2), at three different thresholds of abundance for the rare cell population (5%, 1%, and 0.1%; by column).

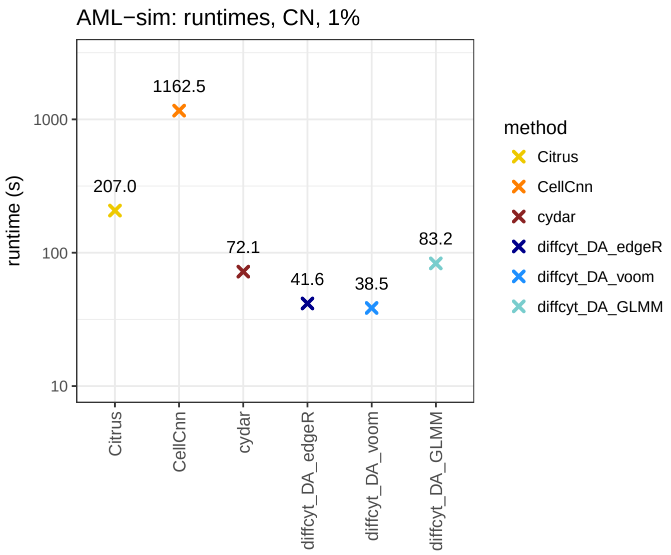

**Supplementary Figure 12. Runtimes: AML-sim, all methods, main simulation.** Runtimes for all methods; testing for differential abundance of cell populations. Runtimes are shown for condition CN vs. healthy at one threshold of abundance for the rare cell population (1%). Text labels indicate runtimes in seconds. All methods were run on a 2014 MacBook Air laptop, 1.7 GHz processor, 8 GB memory, using a single processor core. For `Citrus`, subsampling was used to select a maximum of 5,000 cells per sample; all other methods were run without subsampling. See Supplementary Note 2 for more details on parameters. Runtimes for the other condition (CBF vs. healthy) and the remaining thresholds of abundance for the rare cell population (5% and 0.1%) were approximately similar.

#### 1.2 BCR-XL-sim: Differential states within cell populations

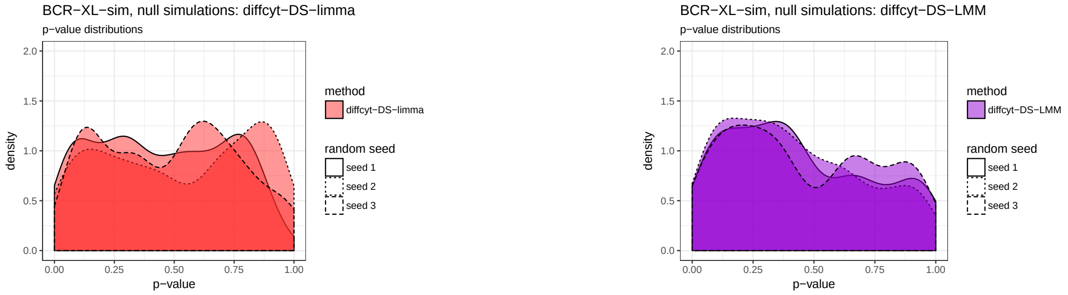

**Supplementary Figure 13. Null simulation p-values: BCR-XL-sim, diffcyt methods.** P-value distributions (densities) for `diffcyt` methods, 3 replicated null simulations; testing for differential states within cell populations. P-value distributions that are approximately uniform across replicates are consistent with data containing no true differential signal.

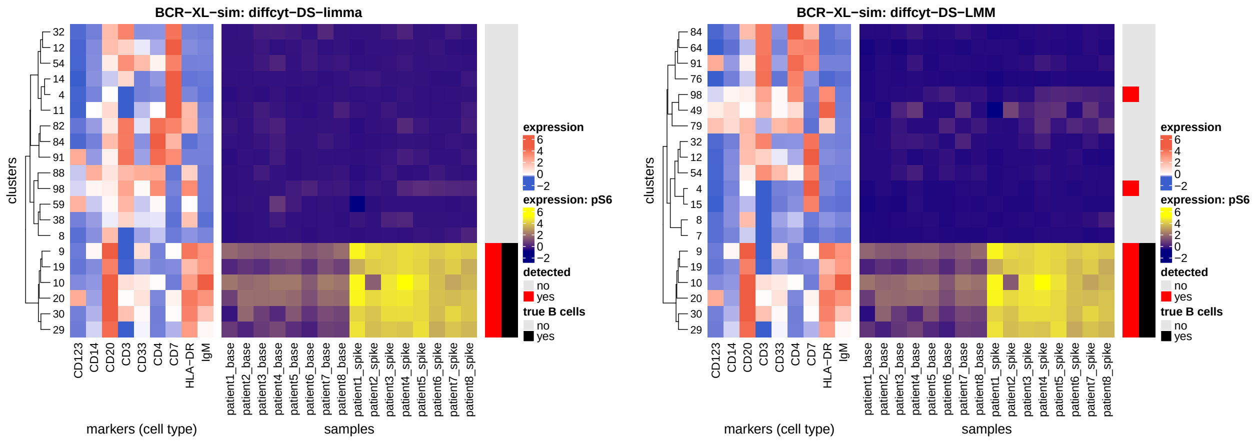

**Supplementary Figure 14. Heatmaps: BCR-XL-sim, diffcyt methods.** Heatmaps showing (i) phenotypes (median arcsinh-transformed expression profiles for cell type markers) (left panel) and (ii) expression of signaling marker pS6 by sample (right panel) for each cluster; testing for differential states within cell populations. Vertical annotation highlights detected significant cluster-marker combinations at 10% FDR (red) and clusters containing >50% true spiked-in cells (black). Color scale for expression of cell type markers is normalized to 1st and 99th percentiles across all clusters and markers. Clusters (rows) are grouped using hierarchical clustering with Euclidean distance and average linkage. Each heatmap shows only the top 20 clusters as ranked by significance levels (for signaling marker pS6; out of 100 clusters total), for easier visibility of the top detected clusters. Panels show results separately for each method.

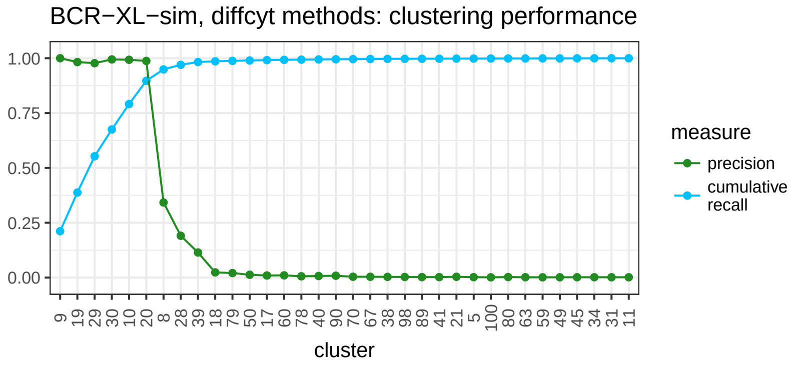

**Supplementary Figure 15. Clustering performance: BCR-XL-sim, diffcyt methods.** Clustering performance measures (precision, cumulative recall) for all clusters containing any true B cells (recall $> 0$); testing for differential states within cell populations. Clusters (horizontal axis) are ordered by recall. Note that the clustering step is the same for both `diffcyt` methods (`diffcyt-DS-limma` and `diffcyt-DS-LMM`).

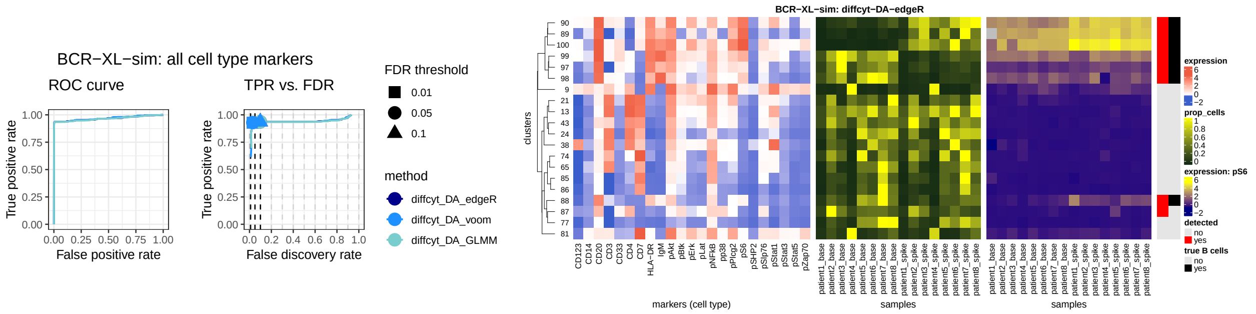

**Supplementary Figure 16. Treating all markers as 'cell type' and testing with DA (instead of DS) methods: BCR-XL-sim; diffcyt methods.** (Left:) Results of performance evaluations for `diffcyt` methods, treating all markers as 'cell type' (i.e. used for clustering); testing for differential abundance of cell populations. Panels show (i) receiver operating characteristic (ROC) curves, and (ii) true positive rate (TPR) vs. false discovery rate (FDR) curves (also showing observed TPR and FDR at FDR cutoffs 1%, 5%, and 10%). (Right:) Heatmap showing (i) phenotypes (median arcsinh-transformed expression profiles for all markers), (ii) relative cluster abundances (proportion of cells per cluster, by sample), and (iii) expression of signaling marker pS6 by sample for each cluster. Vertical annotation highlights detected significant clusters at 10% FDR (red) and clusters containing >50% true spiked-in cells (black). Heatmap is shown for one method only (`diffcyt-DA-edgeR`); heatmaps for the other methods are similar. The group of 6 significant clusters (clusters 89, 90, 97, 98, 99, 100) matches the expected phenotype from the main results (B cells identified by high expression of CD20, with either high or low expression of pS6). However, compared to the main results (heatmaps in Supplementary Figure 16), these results are more difficult to interpret, since cluster phenotypes (expression profiles) may mix elements from canonical 'cell type' and 'cell state' phenotypes.

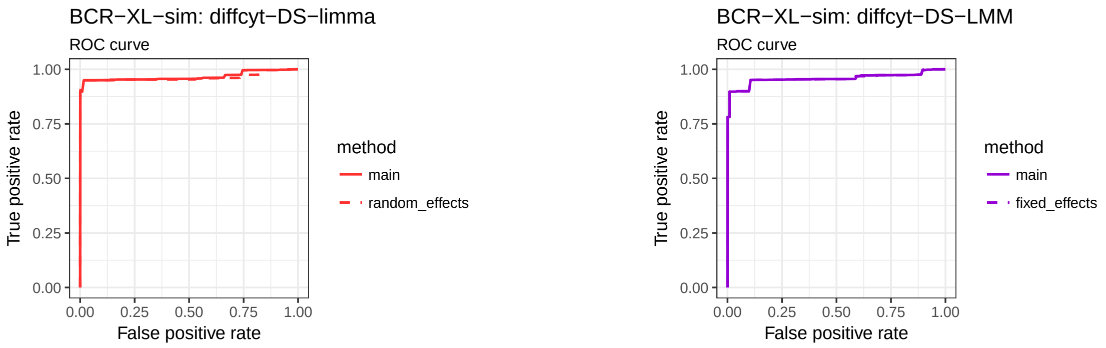

**Supplementary Figure 17. Performance metrics: BCR-XL-sim, diffcyt-DS-limma and diffcyt-DS-LMM, alternative methodologies for patient IDs.** Results of performance evaluations for (i) `diffcyt-DS-limma`, main results vs. using random effects instead of fixed effects for patient IDs (using `limma duplicateCorrelation` methodology) (left panel); and (ii) `diffcyt-DS-LMM`, main results vs. using fixed effects instead of random effects for patient IDs (right panel); testing for differential states within cell populations. Results are displayed using receiver operating characteristic (ROC) curves.

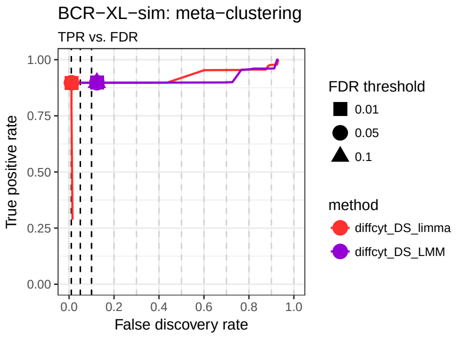

**Supplementary Figure 18. Performance metrics: BCR-XL-sim, diffcyt methods, using FlowSOM meta-clustering.** Results of performance evaluations for `diffcyt` methods, using 20 meta-clusters in FlowSOM clustering algorithm; testing for differential states within cell populations. Results are displayed using true positive rate (TPR) vs. false discovery rate (FDR) curves (also showing observed TPR and FDR at FDR cutoffs 1%, 5%, and 10%).

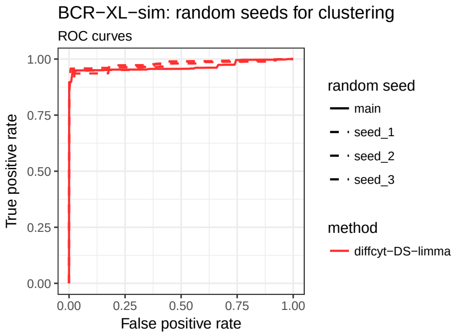

**Supplementary Figure 19. Performance metrics: BCR-XL-sim, diffcyt methods, varying random seeds for clustering.** Results of performance evaluations for `diffcyt-DS-limma`, main results and 3 additional replicates using varying random seeds for clustering step; testing for differential states within cell populations. Results are displayed using receiver operating characteristic (ROC) curves. Results for method `diffcyt-DS-LMM` are approximately similar.

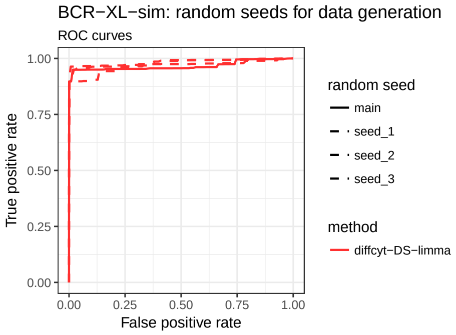

**Supplementary Figure 20. Performance metrics: BCR-XL-sim, diffcyt methods, varying random seeds for data generation.** Results of performance evaluations for `diffcyt-DS-limma`, main results and 3 additional replicates using varying random seeds for data generation; testing for differential states within cell populations. Results are displayed using receiver operating characteristic (ROC) curves. Results for method `diffcyt-DS-LMM` are approximately similar.

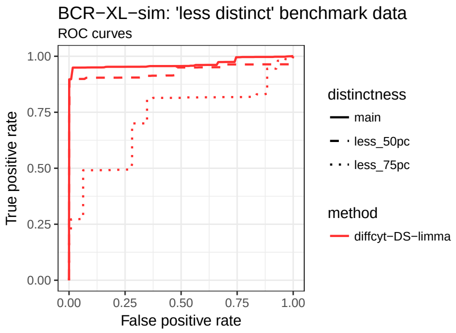

**Supplementary Figure 21. Performance metrics: BCR-XL-sim, diffcyt methods, 'less distinct' benchmark data.** Results of performance evaluations for `diffcyt-DS-limma`, main results and 50% and 75% 'less distinct' benchmark datasets; testing for differential states within cell populations. Results are displayed using receiver operating characteristic (ROC) curves. Results for method `diffcyt-DS-LMM` are approximately similar.

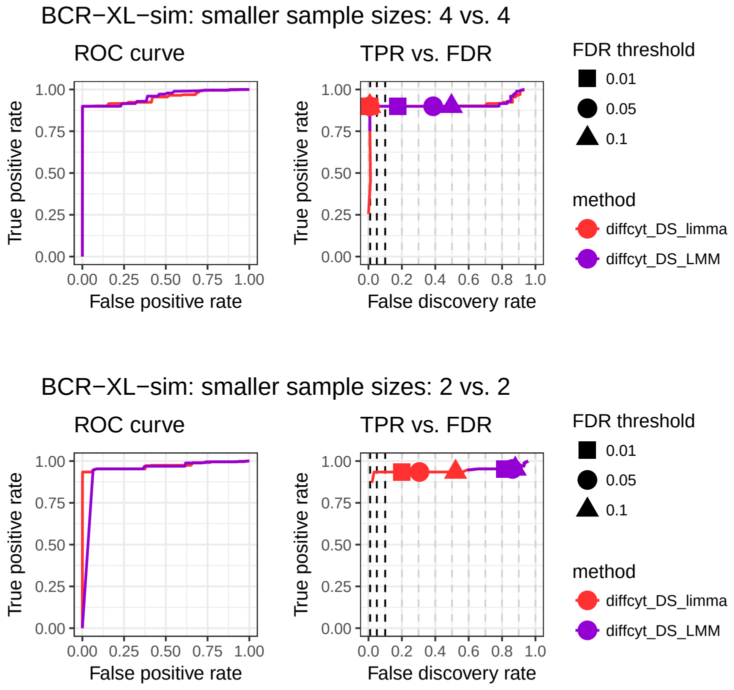

**Supplementary Figure 22. Performance metrics: BCR-XL-sim, diffcyt methods, smaller sample sizes.** Results of performance evaluations for `diffcyt` methods, using subsets of the full number of samples; testing for differential states within cell populations. 4 vs. 4 samples (top) and 2 vs. 2 samples (bottom); the full dataset contains 8 vs. 8 samples. Panels show (i) receiver operating characteristic (ROC) curves, and (ii) true positive rate (TPR) vs. false discovery rate (FDR) curves (also showing observed TPR and FDR at FDR cutoffs 1%, 5%, and 10%).

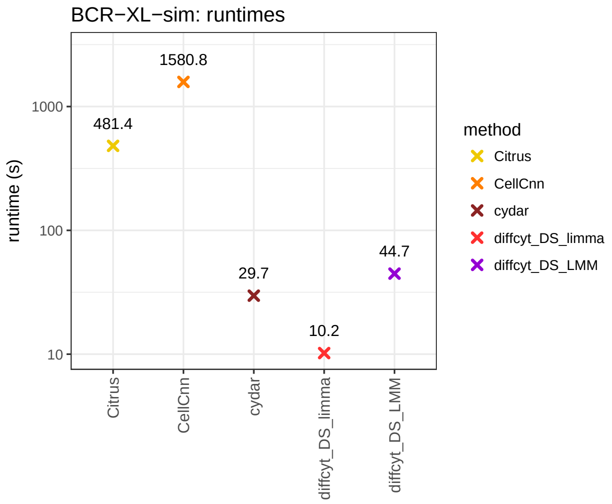

**Supplementary Figure 23. Runtimes: BCR-XL-sim, all methods, main simulation.** Runtimes for all methods; testing for differential states within cell populations. Text labels indicate runtimes in seconds. All methods were run on a 2014 MacBook Air laptop, 1.7 GHz processor, 8 GB memory, using a single processor core. For `Citrus`, subsampling was used to select a maximum of 5,000 cells per sample; all other methods were run without subsampling. `cydar` was run using only a subset of markers ('cell type' markers excluding CD45, plus pS6). See Supplementary Note 2 for more details on parameters.

#### 1.3 Anti-PD-1: Re-analysis of experimental data

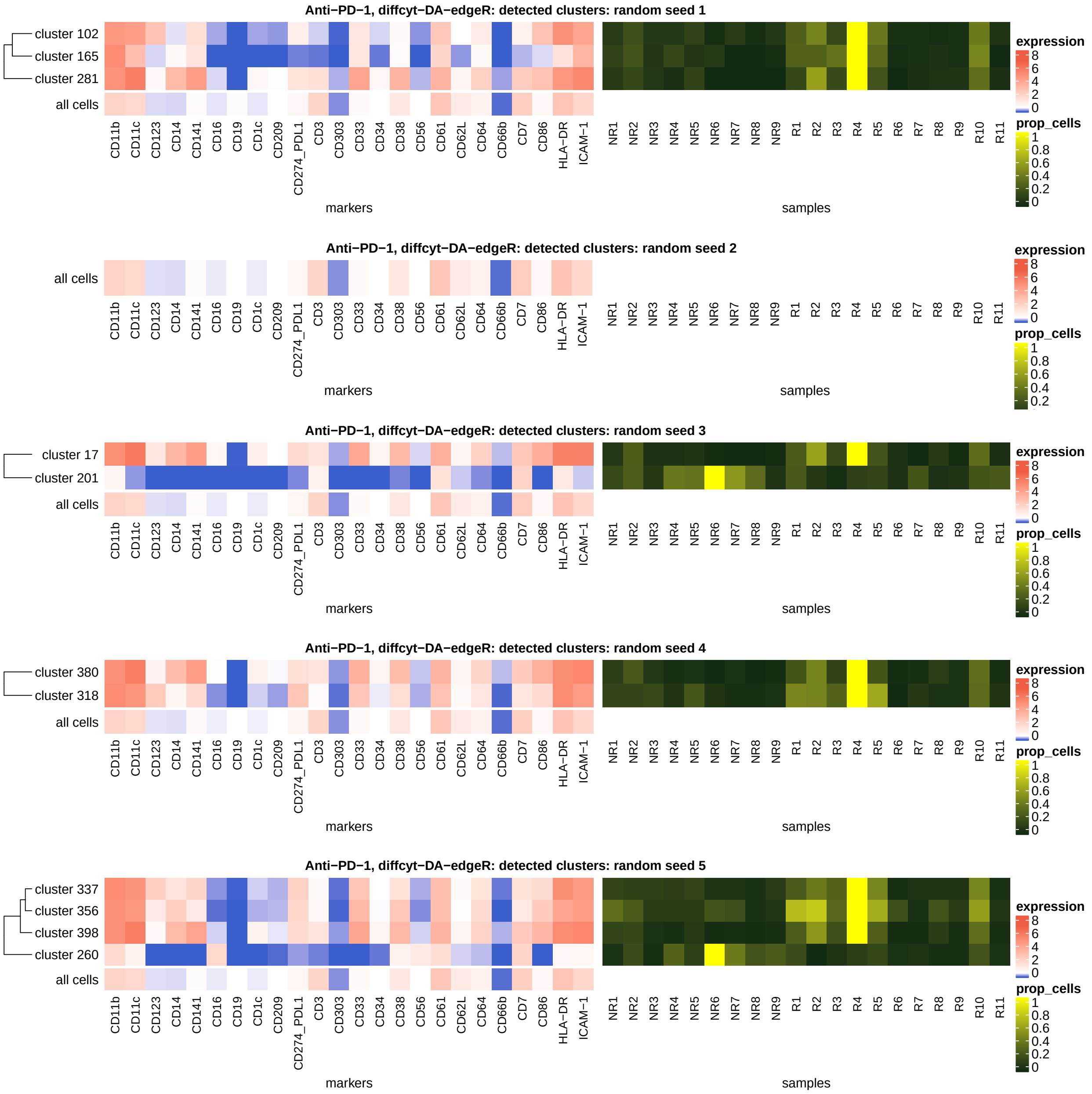

**Supplementary Figure 24. Sensitivity to random seeds: Anti-PD-1, diffcyt-DA-edgeR.** Results for 5 additional runs using different random seeds (for re-analysis of experimental dataset Anti-PD-1 using `diffcyt-DA-edgeR`; testing for differential abundance of cell populations between baseline samples from responder and non-responder groups of patients). Between 0 and 4 significant clusters are detected per run. Clusters matching the phenotype of interest are detected in 4 out of 5 runs (additional possible false positives are represented by cluster 201 for random seed 3 and cluster 260 for random seed 5; see previous figure and main text). Heatmaps show (i) phenotype (median arcsinh-transformed marker expression profiles) of significant detected clusters at 10% false discovery rate (FDR), compared to all cells (left panel); and (ii) relative cluster abundances (proportion of cells per cluster, by sample) (right panel) for the detected clusters. Heatmap rows (clusters) are grouped by hierarchical clustering with Euclidean distance and average linkage. NR = non-responders, R = responders.

#### 1.4 BCR-XL: Re-analysis of experimental data

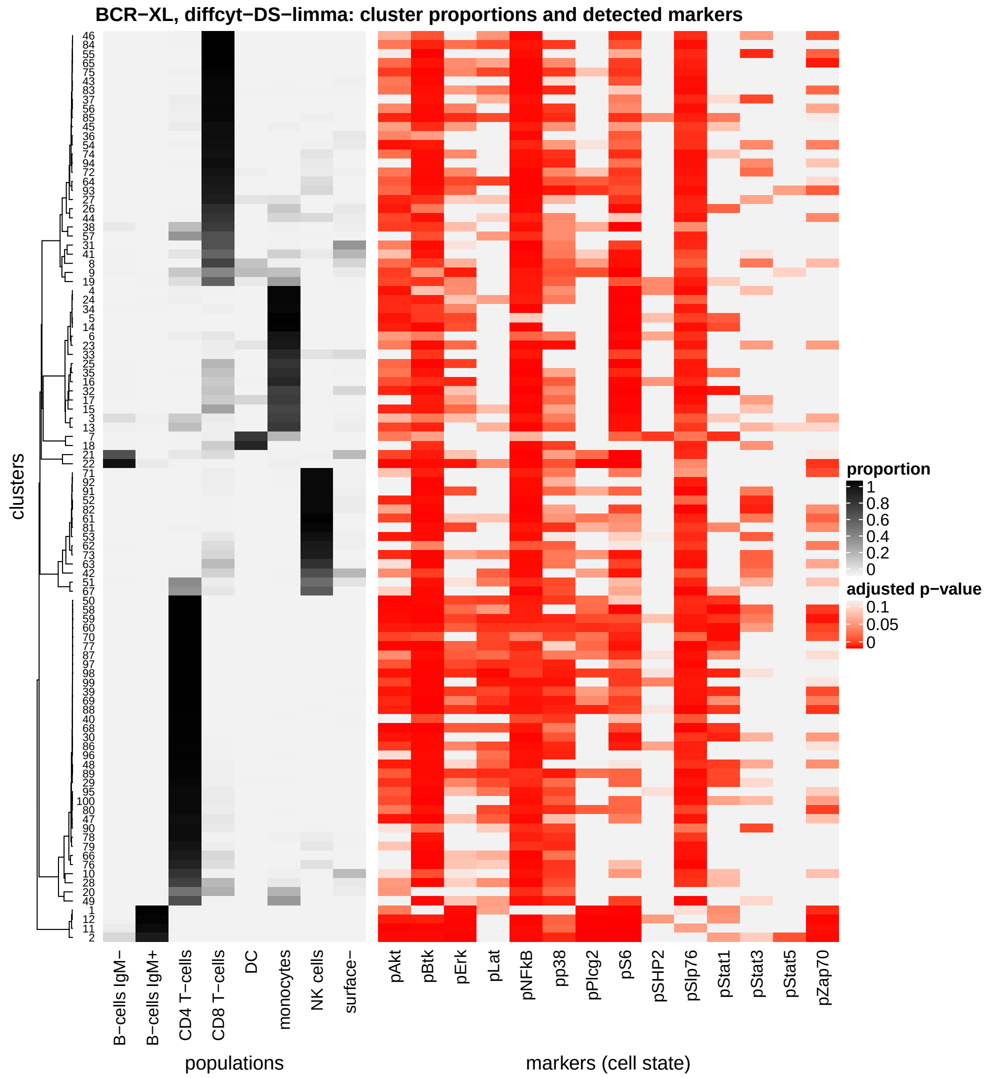

**Supplementary Figure 25. Main results: BCR-XL, diffcyt-DS-limma.** Results for re-analysis of experimental dataset BCR-XL using `diffcyt-DS-limma`; testing for differential states within cell populations. Heatmap shows (i) proportions of cells per cluster matching to each true population (left panel); and (ii) adjusted p-values from the differential tests for each cell state marker for each cluster (right panel). All clusters (total 100 clusters) and cell state markers (14 cell state markers) are shown. Rows (clusters) are grouped by hierarchical clustering with Euclidean distance and average linkage (on the proportions). True (reference) cell population labels are sourced from [1] (see Supplementary Note 1).

### Supplementary Note 1: Benchmark datasets

#### 2.1 AML-sim

##### 2.1.1 Summary

The 'AML-sim' dataset is a semi-simulated dataset designed to evaluate the performance of methods for detecting differential abundance of a single rare cell population.

The dataset consists of several samples of bone marrow mononuclear cells (BMMCs) from healthy individuals, with small percentages of acute myeloid leukemia (AML) blast cells computationally 'spiked-in' at various thresholds of abundance. This simulates the phenotype of minimal residual disease (MRD) in AML patients. The question of interest is to detect the differentially abundant rare population of AML blast cells, even at extremely small thresholds.

The data generation concept and strategy are based on a similar benchmark dataset created by [2], who used their dataset to demonstrate the performance of `CellCnn`. The original data is sourced from [3], and is available from Cytobank at the following links. Gating plots for the blast cells are also shown in [3], Supplemental Data S3B.

- all cells (also contains gating scheme for $CD34^{+}CD45^{mid}$ cells, i.e. blasts): https://community.cytobank.org/cytobank/experiments/46098/illustrations/121588
- blasts (repository cloned from the one for 'all cells' above, using the gating scheme for $CD34^{+}CD45^{mid}$ cells; this allows `.fcs` files for the subset to be exported): https://community.cytobank.org/cytobank/experiments/63534/illustrations/125318

##### 2.1.2 Details on data generation strategy

The original dataset [3] consists of 5 healthy samples and 16 AML samples. Several of the AML samples have been classified into two subtypes [2]: 'cytogenetically normal' or CN (patients SJ10, SJ12, and SJ13), and 'core-binding factor translocation' or CBF (patients SJ1, SJ2, SJ3, SJ4, and SJ5).

For the AML-sim dataset, we use all 5 healthy samples (labeled H1–H5), one CN sample, and one CBF sample. The CN and CBF samples correspond to individuals SJ10 (CN) and SJ4 (CBF) in the meta-data spreadsheets. (Note that the `.fcs` filenames do not correspond to the correct sample names from the meta-data spreadsheet; see comments in data preparation script `prepare_data_AML_sim_main.R` at https://github.com/lmweber/diffcyt-evaluations for more details).

To generate the AML-sim dataset, we split each healthy sample (H1–H5) into 3 equal parts. The first part is kept as the 'healthy' sample. In the second part, we computationally 'spike in' small percentages of randomly selected blast cells from the CN sample (SJ10). Similarly, in the third part, we spike in small percentages of randomly selected blast cells from the CBF sample (SJ4). The blast cells are spiked-in at three different thresholds (5%, 1%, and 0.1%) of the number of healthy cells for each sample, to create several datasets with varying levels of 'rareness' for the population of interest. Different random seeds are used for splitting each healthy sample and for selecting blast cells for each sample and threshold, to ensure that the sets of cells are not identical across replicates. The objective is then to detect the differentially abundant population of CN blasts in a comparison of the 5 healthy samples versus 5 'healthy + CN' samples at each threshold; and similarly for the CBF blasts.

Supplementary Table 1 shows the numbers of cells for each healthy sample (H1–H5) and threshold, and Supplementary Table 2 lists the protein markers in this dataset. There are 31 protein markers in total, including 16 surface markers used to define cell types, and 15 intracellular markers used to characterize cell states. For the AML-sim dataset, we only require the surface markers, since we are only interested in detecting differential abundance of cell populations (i.e. cell types). Supplementary Figure 26 (left panel) displays marker expression profiles (distributions of arcsinh-transformed expression values for each protein marker) for blast cells in each condition: healthy, CN, and CBF.

| Sample | Total no. cells | 5% | 1% | 0.1% |
|---|---|---|---|---|
| H1 | 15,394 | 257 | 52 | 6 |
| H2 | 26,633 | 444 | 89 | 9 |
| H3 | 21,246 | 355 | 71 | 8 |
| H4 | 41,848 | 698 | 140 | 14 |
| H5 | 52,472 | 875 | 175 | 18 |

**Supplementary Table 1. AML-sim dataset: number of cells.** For each healthy sample (H1–H5), the columns show the total number of cells, and the numbers of 'spiked-in' cells randomly selected from either the CN or CBF sample to create the simulated samples. The healthy samples are split into 3 equal parts; CN or CBF cells are then spiked-in at several thresholds (5%, 1%, or 0.1%) of the number of cells in the corresponding 'healthy' sample (which consists of one third of the total cells for that sample); for example, H1: $5\% * 15394/3 = 257$ cells.

| Protein marker class | Protein marker names |
|---|---|
| Cell type | CD117, CD11b, CD123, CD15, CD19, CD3, CD33, CD34, CD38, CD41, CD44, CD45, CD47, CD64, CD7, HLA-DR |
| Cell state | cCaspase3, p4EBP1, pAKT, pAMPK, pc-Cbl, pCREB, pErk1-2, pP38, pPLCg2, pRb, pS6, pSTAT1, pSTAT3, pSTAT5, pZap70-Syk |

**Supplementary Table 2. AML-sim dataset: protein markers.** Summary of protein marker classes (cell type or cell state) and names in the AML-sim dataset. There are 31 protein markers in this dataset, including 16 surface markers used to define cell types, and 15 intracellular markers representing cell states. For the AML-sim dataset, we require only the surface markers.

##### 2.1.3 Randomized benchmark datasets

Several steps in the data generation strategy described above depend on random sampling of cells. To investigate the sensitivity to this random sampling, we generated several additional replicates of the AML-sim dataset using different random seeds.

The modified random seeds affected the random sampling for (i) splitting each healthy sample (H1–H5) into three equal parts; and (ii) selecting subsets of blast cells from the CN and CBF samples at each spike-in threshold (5%, 1%, and 0.1%). We generated three randomized replicates of the AML-sim dataset using this strategy.

##### 2.1.4 'Less distinct' populations of interest

In the main AML-sim dataset, the rare populations of CN and CBF blast cells have marker expression profiles that are clearly distinct from healthy blasts (Supplementary Figure 26, left panel). This arguably presents a relatively 'easy' task for the methods used to test for differential abundance.

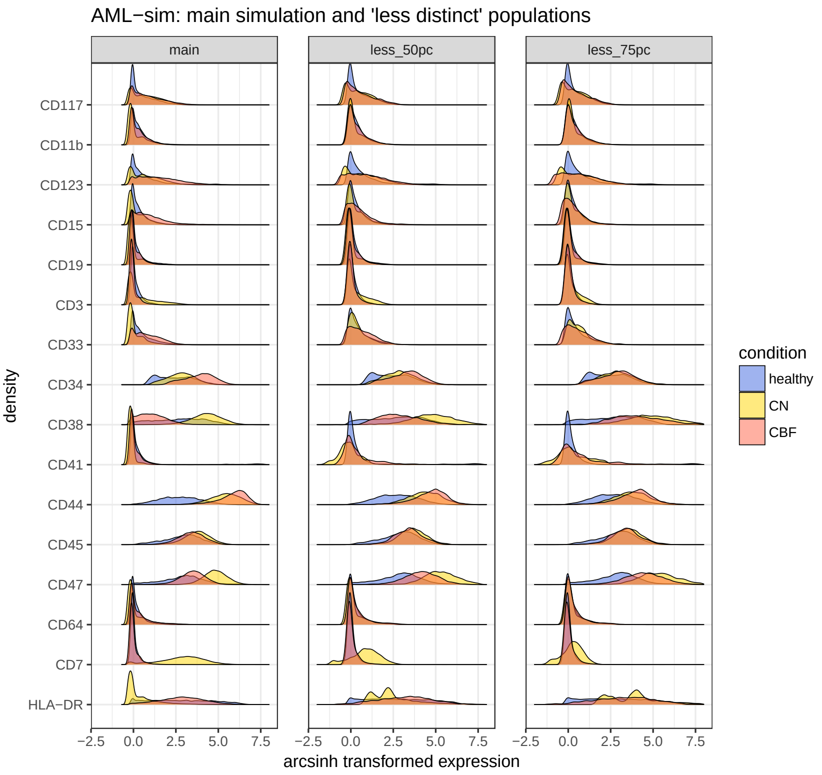

**Supplementary Figure 26. AML-sim dataset: marker distributions for blasts cells, main simulation and 'less distinct' simulations.** Distributions of arcsinh-transformed expression values for each protein marker (marker expression profiles) for blast cells, by condition, for the main simulation (left panel) and the 50% and 75% 'less distinct' simulations (middle and right panels). Healthy samples (H1–H5) are combined into a single curve.

In order to further evaluate the sensitivity of the methods, we generated additional simulations where the CN and CBF blast cell populations have been modified to be 'less distinct' from the healthy blasts, according to their marker expression profiles. This increases the level of difficulty for the differential testing methods.

We created 'less distinct' AML blast cell populations (CN and CBF) by scaling the marker expression profiles of these populations, reducing the differences in both median and standard deviation of the arcsinh-transformed expression profiles by certain proportions, compared to healthy blasts. We created two new simulated datasets: reducing the differences in median and standard deviation by 50% and 75%. (In other words, for each cell in the AML blast cell population of interest, 50% or 75% of the difference in medians between AML blasts and healthy blasts was subtracted, followed by dividing the values to reduce the difference in standard deviations by 50% or 75%). Supplementary Figure 26 (middle and right panels) displays the resulting marker expression profiles.

##### 2.1.5 Null simulations

In order to investigate the error rates of the methods for testing for differential abundance, we also generated 'null' simulations based on the AML-sim dataset. The null simulations do not include any true 'spiked-in' cells; therefore, any detected signals (significant differentially abundant clusters) will be false positives. We generated the null simulations by splitting each healthy sample (H1–H5) into two equal parts, and then tested for differential abundance of cell populations between these two parts. This was repeated three times using different random seeds, to generate three replicates.

#### 2.2 BCR-XL-sim

##### 2.2.1 Summary

The 'BCR-XL-sim' dataset is a semi-simulated dataset designed to evaluate the performance of methods for detecting differential states within cell populations.

The dataset consists of two groups of paired samples of healthy peripheral blood mononuclear cells (PBMCs), where one group contains a computationally 'spiked-in' population of B cells from matched samples stimulated with B cell receptor / Fc receptor cross-linker (BCR-XL). The stimulated cells are from the same individual as the healthy cells for each pair, preserving the paired data structure. The stimulated B cells contain a known signal of elevated expression of several signaling markers; the strongest signal is for phosphorylated ribosomal protein S6 (pS6), as previously described by [4] and [5]. The aim is to detect differential expression of the signaling state marker pS6 in B cells between the two groups.

The original dataset is from [5], and has previously been used for benchmarking evaluations by [4] (who used it to demonstrate the performance of `Citrus`) and [1].

- The original data is available from Cytobank: https://community.cytobank.org/cytobank/experiments/15713/download_files
- Additional information is available from the `Citrus` wiki page: https://github.com/nolanlab/citrus/wiki/PBMC-Example-1

Cell population labels (required to identify B cells) are reproduced from [1], where they were generated using a strategy of expert-guided manual merging of automatically generated clusters from the `FlowSOM` algorithm.

##### 2.2.2 Details on data generation strategy

The original dataset [5] consisted of samples of healthy PBMCs from 8 individuals, where samples from each individual were stimulated with a number of different signaling inhibitors in order to investigate properties of cell signaling networks. For the BCR-XL-sim dataset, we use samples from the unstimulated reference condition and samples stimulated with B cell receptor / Fc receptor cross-linker (BCR-XL). Therefore, we have 16 original samples, in an 8 vs. 8 paired design.

As previously described [1, 4, 5], this dataset contains strong differential expression signals for several signaling state markers in several cell populations. In particular, one of the strongest effects is differential expression of phosphorylated S6 (pS6) in B cells (see [1], Figure 27).

We construct the 'BCR-XL-sim' benchmark dataset as follows. First, we select the unstimulated reference sample from each pair, and randomly split this into two halves. Then, in one half, we replace the B cells with an equivalent number of B cells from the corresponding paired sample from the BCR-XL stimulated condition. This introduces a differential expression signal in B cells for several signaling state markers, including pS6. Methods are then evaluated by their ability to detect differential expression of pS6 in B cells between the two conditions.

Supplementary Table 3 summarizes the number of cells in this dataset, and Supplementary Table 4 lists the protein markers. There are 24 protein markers in total, including 10 surface markers (9 of which are used to define cell types), and 14 intracellular markers used to characterize cell states. Note that CD45 is excluded from the set of surface markers used to define cell types by clustering, since almost all cells in this dataset have high expression of CD45; hence CD45 is not informative for distinguishing cell populations. Supplementary Figure 27 (left panel) displays the marker expression profiles (distributions of arcsinh-transformed expression values for each protein marker) in each condition.

| Individual | Total cells | B cells |
|---|---|---|
| 1 | 2,739 | 184 |
| 2 | 16,725 | 686 |
| 3 | 9,434 | 1,091 |
| 4 | 6,906 | 422 |
| 5 | 11,962 | 830 |
| 6 | 11,038 | 885 |
| 7 | 15,974 | 1,139 |
| 8 | 13,670 | 821 |

**Supplementary Table 3. BCR-XL-sim dataset: number of cells.** For each individual, the columns show the total number of cells and the number of B cells in the unstimulated (reference) sample (B cells are also included in the total). For the BCR-XL-sim dataset, each unstimulated sample is split into two equal parts, and the B cells in one part are replaced with an equivalent number of B cells from the corresponding paired sample from the stimulated (BCR-XL) condition.

| Protein marker class | Protein marker names |
|---|---|
| Cell type | CD123, CD14, CD20, CD3, CD33, CD4, CD45, CD7, HLA-DR, IgM |
| Cell state | pAkt, pBtk, pErk, pLat, pNFkB, pp38, pPlcg2, pS6, pSHP2, pSlp76, pStat1, pStat3, pStat5, pZap70 |

**Supplementary Table 4. BCR-XL-sim dataset: protein markers.** Summary of protein marker classes (cell type or cell state) and names in the BCR-XL-sim dataset. There are 24 protein markers in this dataset, including 10 surface markers (9 of which are used to define cell types), and 14 intracellular markers representing cell states. Note that CD45 is excluded from the set of surface markers used to define cell types by clustering, since almost all cells in this dataset have high expression of CD45; hence CD45 is not informative for distinguishing cell populations.

##### 2.2.3 Randomized benchmark datasets

Several steps in the data generation strategy described above depend on random sampling of cells. To investigate the sensitivity to this random sampling, we generated several additional replicates of the BCR-XL-sim dataset using different random seeds.

The modified random seeds affected the random sampling for (i) splitting each unstimulated (reference) sample into two equal parts; and (ii) selecting B cells from the stimulated (BCR-XL) condition to use as 'spiked-in' cells. We generated three randomized replicates of the BCR-XL-sim dataset using this strategy.

##### 2.2.4 'Less distinct' populations of interest

In the main BCR-XL-sim dataset, the spiked-in populations of stimulated B cells have marker expression profiles that are clearly distinct from healthy B cells (Supplementary Figure 27, left panel). This arguably presents a relatively 'easy' task for the methods used to test for differential states within cell populations.

In order to further evaluate the sensitivity of the methods, we generated additional simulations where the stimulated B cells have been modified to be 'less distinct' from the

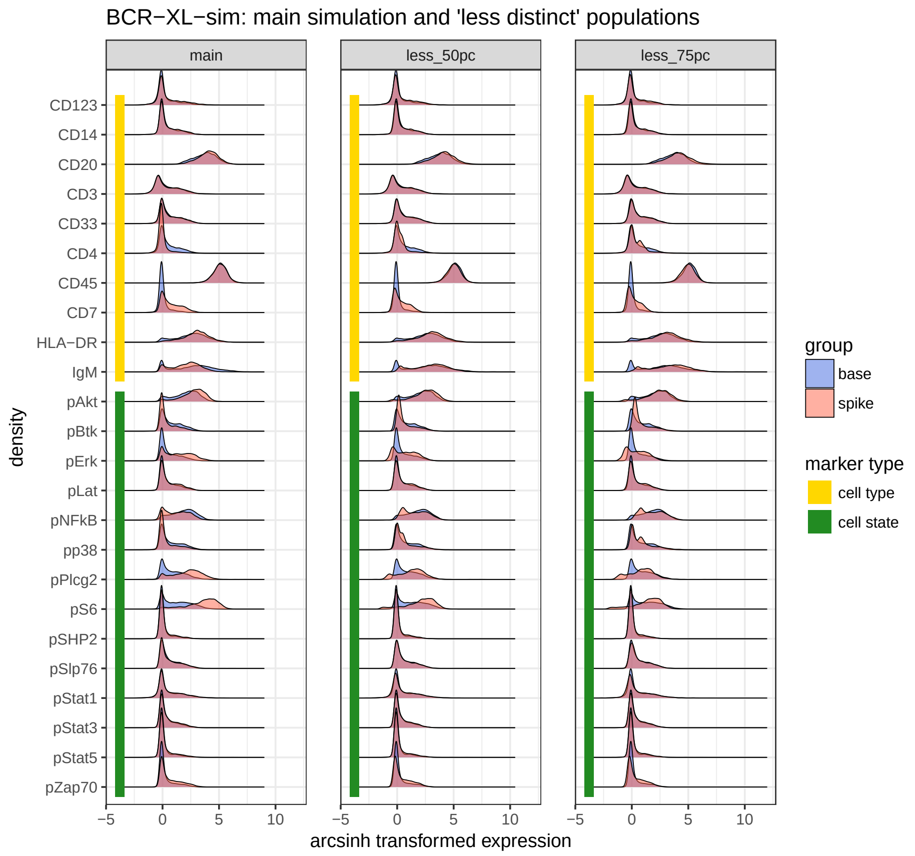

**Supplementary Figure 27. BCR-XL-sim dataset: marker distributions for B cells, main simulation and 'less distinct' simulations.** Distributions of arcsinh-transformed expression values for each protein marker (marker expression profiles) for B cells, by condition, for the main simulation and the 50% and 75% 'less distinct' simulations.

healthy B cells, according to their marker expression profiles. This increases the level of difficulty for the differential testing methods.

We created 'less distinct' stimulated B cell populations by scaling the marker expression profiles of these populations, reducing the differences in both median and standard deviation of the arcsinh-transformed expression profiles by certain proportions, compared to healthy B cells. We created two new simulated datasets: reducing the differences in median and standard deviation by 50% and 75%. (In other words, for each cell in the stimulated B cell population, 50% or 75% of the difference in medians between stimulated and healthy B cells was subtracted, followed by dividing the values to reduce the difference in standard deviations by 50% or 75%). Supplementary Figure 27 (middle and right panels) displays the resulting marker expression profiles.

##### 2.2.5 Null simulations

In order to investigate the error rates of the methods for testing for differential states within cell populations, we also generated 'null' simulations based on the BCR-XL-sim dataset. The null simulations do not include any true 'spiked-in' cells; therefore, any detected signals (significant differential expression of cell state markers within clusters) will be false positives. We generated the null simulations by splitting each unstimulated (reference) sample into two equal parts, and then tested for differential states within cell populations between these two parts. This was repeated three times using different random seeds, to generate three replicates.

#### 2.3 Anti-PD-1

The 'Anti-PD-1' dataset is an experimental dataset from a recent study using mass cytometry to characterize immune cell subsets in peripheral blood from melanoma skin cancer patients treated with anti-PD-1 immunotherapy [6].

One of the key results of this study was that the frequency of CD14$^+$ CD16$^-$ HLA-DR$^{hi}$ monocytes in baseline samples (taken from patients prior to treatment) was a strong predictor of survival in response to immunotherapy treatment. In particular, analysis using `CellCnn` [2] detected a small subpopulation of CD14$^+$ CD33$^+$ HLA-DR$^{hi}$ ICAM-1$^+$ CD64$^+$ CD141$^+$ CD86$^+$ CD11c$^+$ CD38$^+$ PD-L1$^+$ CD11b$^+$ monocytes within this population; the frequency of this subpopulation in baseline samples was strongly associated with responder status following immunotherapy treatment. (See section 'Identification of a monocyte signature using CellCnn', referring to results for 'myeloid panel', in [6]).

Here, we re-analyzed this dataset using the `diffcyt` methods, in order to test whether the `diffcyt` methods could reproduce this known result from a published experimental dataset. Note that this dataset contains a strong batch effect, due to sample acquisition on two different days [6].

Since this is an experimental dataset, there is no known 'truth' that can be used to calculate statistical performance metrics (i.e. unlike the simulated datasets). Instead, the results are evaluated qualitatively, using visualizations to determine whether the `diffcyt` methods detect differentially abundant clusters corresponding to the previously validated differentially abundant subpopulation of monocytes.

#### 2.4 BCR-XL

The 'BCR-XL' dataset refers to the original experimental data from [5], which was also used to construct the BCR-XL-sim semi-simulated dataset described above (see section 'BCR-XL-sim' for more details about the original dataset).

As described above, the BCR-XL dataset contains known strong differential signals for several signaling state markers in several cell populations; one of the strongest signals is for differential expression of pS6 in B cells. Here, we applied the `diffcyt` methods directly to the unmodified original dataset (BCR-XL stimulation condition only), in order to test whether the `diffcyt` methods could detect the known strong differential signals.

As for the Anti-PD-1 dataset, this is an experimental dataset, which does not contain a known 'truth' that can be used to calculate statistical performance metrics. Therefore, the results are evaluated qualitatively using visualizations. (However, reference cell population labels are available from [1], as described above for the BCR-XL-sim dataset; these can be used to generate more informative visualizations.)

#### 2.5 Data and code availability

Data files for all benchmark datasets are available in `.fcs` format from FlowRepository (repository ID: FR-FCM-ZYL8) at http://flowrepository.org/id/FR-FCM-ZYL8. The benchmark datasets can also be accessed in `SummarizedExperiment` and `flowSet` Bioconductor object formats through the `HDCytoData` Bioconductor package, available at http://bioconductor.org/packages/HDCytoData.

R code scripts to reproduce all data preparation and simulation steps and generate all figures are available from GitHub at https://github.com/lmweber/diffcyt-evaluations.

### Supplementary Note 2:
### Comparisons with existing methods

#### 3.1 Citrus

##### Overview

`Citrus` [4] identifies cell populations associated with a clinical endpoint such as disease status by: (i) generating a hierarchy of clusters and calculating feature values such as cluster abundance or median functional marker expression for every cluster in the hierarchy, and (ii) fitting a regularized classification or regression model, which automatically selects 'stratifying' features associated with the endpoint of interest.

The `Citrus` results consist of a list of 'differential features' at the cluster level, e.g. clusters that have been detected as differentially abundant, or clusters with differential expression of functional markers. To compare performance with other methods, we require unique results at the cell level; we achieve this by assigning the cluster-level differential status to all cells within each detected cluster. Since `Citrus` does not return any continuous-valued scores or p-values, it is not possible to rank clusters (or cells) by their differential evidence: clusters (or cells) are either selected as differential or not. Therefore, receiver operating characteristic (ROC) curves and true positive rate (TPR) vs. false discovery rate (FDR) curves consist of straight-line segments, and it is not possible to calculate the observed TPR, FDR, and false positive rate (FPR) at specific FDR cutoffs.

##### Availability

`Citrus` is available as an R package for download from GitHub (https://github.com/nolanlab/citrus/). Installation instructions are provided on the `Citrus` 'wiki', accessible via the GitHub page. (However, installation on our Mac system required several additional steps: customized compiler setup using 'clang4' and 'libomp', customization of R 'Makevars', and installation of packages from source. These steps are partially documented on the wiki page. Alternatively, `Citrus` can be installed more easily on Linux systems.) The R package also includes a graphical user interface; and `Citrus` is also available within the Cytobank commercial online analysis platform.

##### Version

We used `Citrus` version 0.08 (the latest version available as of 4 May 2018); with R version 3.5.0.

##### Parameter settings

We ran `Citrus` on the semi-simulated benchmark datasets (see Supplementary Note 1) using the following parameter settings:

###### *AML-sim dataset*

- Model type: `family = "classification"; modelTypes = "glmnet"; nFolds = 1`
- Feature type: `featureType = "abundances"`
- Columns: using 'cell type' markers for clustering

- Maximum number of cells per sample: `fileSampleSize = 5000`
- Minimum cluster size: `minimumClusterSizePercent = 0.001` (i.e. 0.1%)
- Transformation and scaling: no additional transformation or scaling. We apply an `arcsinh` transform with `cofactor = 5` separately prior to analysis.
- Number of processor cores: `n_cores = 1`
- Differential features: using `cv.min` results (this tends to return a larger set of differential features, since the regularization threshold is less stringent than for the alternative option `cv.1se`)
- Experimental design and setup for multiple contrasts: run `Citrus` pipeline separately for each condition versus healthy (CN vs. healthy, CBF vs. healthy)

***BCR-XL-sim dataset***

- Model type: `family = "classification"; modelTypes = "glmnet"; nFolds = 1`
- Feature type: `featureType = "medians"`
- Columns: using 'cell type' markers (excluding CD45) for clustering, and 'cell state' (functional) markers for features (medians)
- Maximum number of cells per sample: `fileSampleSize = 5000`
- Minimum cluster size: `minimumClusterSizePercent = 0.01` (i.e. 1%)
- Transformation and scaling: no additional transformation or scaling. We apply an `arcsinh` transform with `cofactor = 5` separately prior to analysis.
- Number of processor cores: `n_cores = 1`
- Differential features: using `cv.min` results (this tends to return a larger set of differential features, since the regularization threshold is less stringent than for the alternative option `cv.1se`)

#### 3.2 CellCnn

##### Overview

CellCnn [2] applies convolutional neural networks in a representation learning approach to detect rare cell populations associated with a condition of interest, such as disease status. Unlike other existing methods, CellCnn is specifically designed for the detection of rare cell populations, which are often of particular biological interest. Note that CellCnn does not distinguish between 'cell type' and 'cell state' markers, since the representation learning approach does not require clusters to be defined explicitly; differential states within cell populations are instead detected as differentially abundant cell populations in the full high-dimensional space.

The CellCnn results consist of continuous 'scores' at the cell level, indicating the likelihood of each cell belonging to each selected 'filter' (detected differential population). To compare performance with other methods, we require unique results at the cell level. Therefore, if multiple filters are selected, we sum the scores to give a single total score per cell. These scores are then used to rank cells by their differential evidence, allowing receiver operating characteristic (ROC) curves and true positive rate (TPR) vs. false discovery rate (FDR) curves to be calculated. However, the scores cannot be interpreted as p-values (in particular, they are not bounded between 0 and 1), so it is not possible to calculate the observed TPR, FDR, and false positive rate (FPR) at specific FDR cutoffs.

##### Availability

CellCnn is available as a Python 2.7 package for download from GitHub (https://github.com/eiriniar/CellCnn). Installation instructions are provided on the GitHub page. The installation procedure includes automated installation of several dependency packages. Users are also required to install a modified version of one dependency package (`fcm`); this is described on the GitHub page. (To run CellCnn on our Linux system, we also needed to make some customized edits to the source code: add lines `import matplotlib` and `matplotlib.use("agg")` to the beginning of the script `plotting.py`.)

##### Version

We used the latest version of CellCnn available from GitHub as of 4 May 2018 (commit `eee86425c7275c7a3763cdea2f5ffb3b3f71549b`, 22 March 2018); with Python version 2.7.14.

##### Parameter settings

We ran CellCnn on the semi-simulated benchmark datasets (see Supplementary Note 1) using the following parameter settings:

###### *AML-sim dataset*

- Markers: CellCnn did not work correctly when using 'cell type' markers only; for some thresholds and conditions, errors were returned, and the CellCnn runs did not complete. For the main results, we used all markers instead (including 'cell state' markers, which are not relevant for defining cell populations). This stabilized the CellCnn runs, and returned results for all thresholds and conditions. (Note that CellCnn does not distinguish between 'cell type' and 'cell state' markers, since the representation learning approach used by `CellCnn` does not require clusters to be defined explicitly. Therefore, including additional, possibly irrelevant markers should generally not seriously affect performance.)

- Transformation: Do not apply separate `arcsinh` transform prior to analysis, since `CellCnn` automatically applies an internal `arcsinh` transform. (An option `--no_arcsinh` is available to disable the internal transform, but this gave errors on our datasets.)
- Size of each training set (number of cells): `--ncell 300` (increased from default of 200 to increase probability that small populations are included in training sets)
- Threshold for choosing responding cell population: `--filter_response_thres 0.3`
- Subset selection: Use option `--subset_selection outlier` for datasets with extremely rare populations (threshold 0.1%). This biases the subsampling for the training sets towards outliers, increasing the probability that rare populations are included.
- Experimental design and setup for multiple contrasts: run `CellCnn` pipeline separately for each condition versus healthy (CN vs. healthy, CBF vs. healthy)

##### *BCR-XL-sim dataset*

- Markers: using all markers excluding CD45 (note that `CellCnn` does not distinguish between 'cell type' and 'cell state' markers)
- Transformation: Do not apply separate `arcsinh` transform prior to analysis, since `CellCnn` automatically applies an internal `arcsinh` transform. (An option `--no_arcsinh` is available to disable the internal transform, but this gave errors on our datasets.)
- Size of each training set (number of cells): `--ncell 300` (increased from default of 200 to increase probability that small populations are included in training sets)
- Threshold for choosing responding cell population: `--filter_response_thres 0.3`

#### 3.3 cydar

##### Overview

cydar [7] detects differentially abundant cell populations by assigning cells to overlapping 'hyperspheres' in the high-dimensional space of protein markers, and controls the spatial false discovery rate (FDR) in the high-dimensional space. A key advantage of cydar is that it explicitly controls the error rate (spatial FDR) and returns results as adjusted p-values. Note that cydar does not distinguish between 'cell type' and 'cell state' markers; differential states within populations are instead detected as differentially abundant populations in the full high-dimensional space.

The cydar results consist of adjusted p-values at the hypersphere level. Since the hyperspheres can overlap, there can be multiple p-values for each cell. However, to compare performance with other methods, we require unique results at the cell level. To achieve this, we assign a unique adjusted p-value to each cell by selecting the smallest adjusted p-value for any hypersphere containing that cell.

##### Availability

cydar is available as an R package from Bioconductor (http://bioconductor.org/packages/cydar). Installation from Bioconductor automatically installs all required dependencies.

##### Version

We used cydar version 1.4.0 (the latest version available as of 4 May 2018); with R version 3.5.0.

##### Parameter settings

We ran cydar on the semi-simulated benchmark datasets (see Supplementary Note 1) using the following parameter settings:

###### *AML-sim dataset*

- No subsampling: option `equalize = FALSE` in function `poolCells` to select all cells
- Transformation: `arcsinh` transform with `cofactor = 5`
- Markers: using 'cell type' markers
- Differential testing: using `edgeR` for differential tests (as described in the `cydar` Bioconductor vignette)
- Experimental design and setup for multiple contrasts: preprocessing and model fitting is performed on combined data from all conditions (CN, CBF, and healthy); differential tests are calculated separately for each contrast (CN vs. healthy, CBF vs. healthy)
- Filtering: remove low-abundance hyperspheres with average log counts per million (CPM) below 1 (reduced from the default of 5)
- Significance threshold for adjusted p-values: 10%

***BCR-XL-sim dataset***

- No subsampling: option `equalize = FALSE` in function `poolCells` to select all cells
- Transformation: `arcsinh` transform with `cofactor = 5`
- Markers: `cydar` did not work correctly when using all markers or all markers excluding CD45 (the recommended approach); no clusters were detected as differentially abundant in this case. For the main results, we used a subset of markers only (lineage markers excluding CD45, plus pS6). (Therefore, the results are not strictly comparable with other methods, where all markers excluding CD45 were used.)
- Differential testing: using `edgeR` for differential tests (as described in the `cydar` Bioconductor vignette)
- Filtering: remove low-abundance hyperspheres with average log counts per million (CPM) below 1 (reduced from the default of 5)
- Significance threshold for adjusted p-values: 10%

#### 3.4 Notes on performance evaluations and comparisons

##### Performance metrics

For the main performance evaluations, methods were evaluated by calculating and comparing (i) receiver operating characteristic (ROC) curves, and (ii) true positive rate (TPR) vs. false discovery rate (FDR) curves (also showing the observed TPR and FDR at FDR cutoffs of 1%, 5%, and 10%).

##### Notes

(i) For cell-level evaluation of `diffcyt` methods, cluster-level p-values and adjusted p-values are assigned to all cells within each cluster. See sections 3.1–3.3 for descriptions of the evaluation strategies for the other methods.

(ii) `Citrus`: ROC curves for `Citrus` consist of straight-line segments, since `Citrus` does not return any continuous scores such as p-values, which could be used to rank clusters (results are binary; clusters are either selected or not). Since no p-values are available, it is also not possible to calculate observed TPR and FDR at given FDR cutoffs.

(iii) `CellCnn`: Scores returned by `CellCnn` cannot be interpreted as p-values, so it is not possible to calculate observed TPR and FDR at given FDR cutoffs.

(iv) `cydar`: For dataset BCR-XL-sim, we ran `cydar` using only a subset of markers ('cell type' markers excluding CD45, plus pS6); running on all markers or all markers excluding CD45 did not work correctly (no significant differential hyperspheres were returned in this case). Therefore, results are not strictly comparable with other methods, where all markers excluding CD45 were used (see Supplementary Note 3).

##### Software versions

Results and figures were generated using `diffcyt` version 1.3.0 (available from GitHub at https://github.com/lmweber/diffcyt/releases) and R version 3.5.0. The versions of `Citrus`, `CellCnn`, and `cydar` used are listed in sections 3.1–3.3.

#### 3.5 Software and code availability

The current release version of the `diffcyt` R package is available from Bioconductor at http://bioconductor.org/packages/diffcyt. The development version (which may include additional updates) is available via the `devel` version of Bioconductor, or from GitHub at https://github.com/lmweber/diffcyt/.

Code scripts to reproduce all performance evaluations and comparisons with existing methods, and to generate all figures, are available from GitHub at https://github.com/lmweber/diffcyt-evaluations.

### Supplementary References

[1] Nowicka, M., Krieg, C., Weber, L. M., Hartmann, F. J., Guglietta, S., Becher, B., Levesque, M. P., and Robinson, M. D. (2017). CyTOF workflow: differential discovery in high-throughput high-dimensional cytometry datasets. *F1000Research*, 6(v2):748.

[2] Arvaniti, E. and Claassen, M. (2017). Sensitive detection of rare disease-associated cell subsets via representation learning. *Nature Communications*, 8(14825):1–10.

[3] Levine, J. H., Simonds, E. F., Bendall, S. C., Davis, K. L., Amir, E.-A. D., Tadmor, M. D., Litvin, O., Fienberg, H. G., Jager, A., Zunder, E. R., Finck, R., Gedman, A. L., Radtke, I., Downing, J. R., Pe'er, D., and Nolan, G. P. (2015). Data-driven phenotypic dissection of AML reveals progenitor-like cells that correlate with prognosis. *Cell*, 162:184–197.

[4] Bruggner, R. V., Bodenmiller, B., Dill, D. L., Tibshirani, R. J., and Nolan, G. P. (2014). Automated identification of stratifying signatures in cellular subpopulations. *Proceedings of the National Academy of Sciences of the USA*, pages E2770–E2777.

[5] Bodenmiller, B., Zunder, E. R., Finck, R., Chen, T. J., Savig, E. S., Bruggner, R. V., Simonds, E. F., Bendall, S. C., Sachs, K., Krutzik, P. O., and Nolan, G. P. (2012). Multiplexed mass cytometry profiling of cellular states perturbed by small-molecule regulators. *Nature Biotechnology*, 30(9):858–867.

[6] Krieg, C., Nowicka, M., Guglietta, S., Schindler, S., Hartmann, F. J., Weber, L. M., Dummer, R., Robinson, M. D., Levesque, M. P., and Becher, B. (2018). High-dimensional single-cell analysis predicts response to anti-PD-1 immunotherapy. *Nature Medicine*, 24(2):144–153.

[7] Lun, A. T. L., Richard, A. C., and Marioni, J. C. (2017). Testing for differential abundance in mass cytometry data. *Nature Methods*, 14(7):707–709.
